# Isoform Specific Regulation of Adenylyl Cyclase 5 by Gβγ

**DOI:** 10.1101/2023.05.02.539090

**Authors:** Yu-Chen Yen, Yong Li, Chun-Liang Chen, Thomas Klose, Val J Watts, Carmen W Dessauer, John J. G. Tesmer

## Abstract

The nine different membrane-anchored adenylyl cyclase isoforms (AC1-9) in mammals are stimulated by the heterotrimeric G protein Gα_s_, but their response to Gβγ regulation is isoform-specific. For example, AC5 is conditionally activated by Gβγ. Here, we report cryo-EM structures of ligand-free AC5 in complex with Gβγ and of a dimeric form of AC5 that could be involved in its regulation. Gβγ binds to a coiled-coil domain that links the AC transmembrane region to its catalytic core as well as to a region (C_1b_) that is known to be a hub for isoform-specific regulation. We confirmed the Gβγ interaction with both purified proteins and cell-based assays. The interface with Gβγ involves AC5 residues that are subject to gain-of-function mutations in humans with familial dyskinesia, indicating that the observed interaction is important for motor function. A molecular mechanism wherein Gβγ either prevents dimerization of AC5 or allosterically modulates the coiled-coil domain, and hence the catalytic core, is proposed. Because our mechanistic understanding of how individual AC isoforms are uniquely regulated is limited, studies such as this may provide new avenues for isoform-specific drug development.

## Introduction

Adenylyl cyclase (AC) catalyzes the formation of cyclic adenosine monophosphate (cAMP), a canonical second messenger in G protein-coupled receptor (GPCR) signaling that regulates a myriad of cellular processes. The first AC isoform was discovered in 1989 from a bovine brain library^1^, and nine membrane-associated AC isoforms (mACs, AC1-9) were ultimately identified in mammals. They are all stimulated by the heterotrimeric G protein Gα_s_ and the small molecule forskolin (except for AC9, which is conditionally activated by forskolin^2^). In addition to these common activators, each isoform responds differently to other signaling partners, such as Ca^2+^, Gβγ, and Gα_i/o_^3^. In this sense, mACs are master integrators of extracellular signals converging from GPCRs that couple to different heterotrimeric G proteins.

All mACs feature highly conserved and homologous C_1a_ and C_2a_ domains that constitute at their interface the active site and the binding site for forskolin. Gα_s_ binds to both proteins at the periphery and thereby stabilizes the bindings sites for ATP and forskolin (**Fig.1A**). The N-terminal and non-catalytic cytoplasmic domains (C_1b_ and C_2b_) are not well conserved among mACs but are well known to play roles in isoform-specific regulation^4–6^. The first structure of an active mAC fragment was the crystal structure of the catalytic core from the C_1a_ domain of AC5 and the C_2a_ domain of AC2 in complex with Gα_s_·GTPγS and forskolin^7^. This structure resolved how Gα_s_·GTPγS and forskolin stabilize the catalytic domain and was the only structural model of a mammalian mAC for over two decades, highlighting the difficulties in studying the structure of intact mACs from higher organisms.

**Figure 1.**
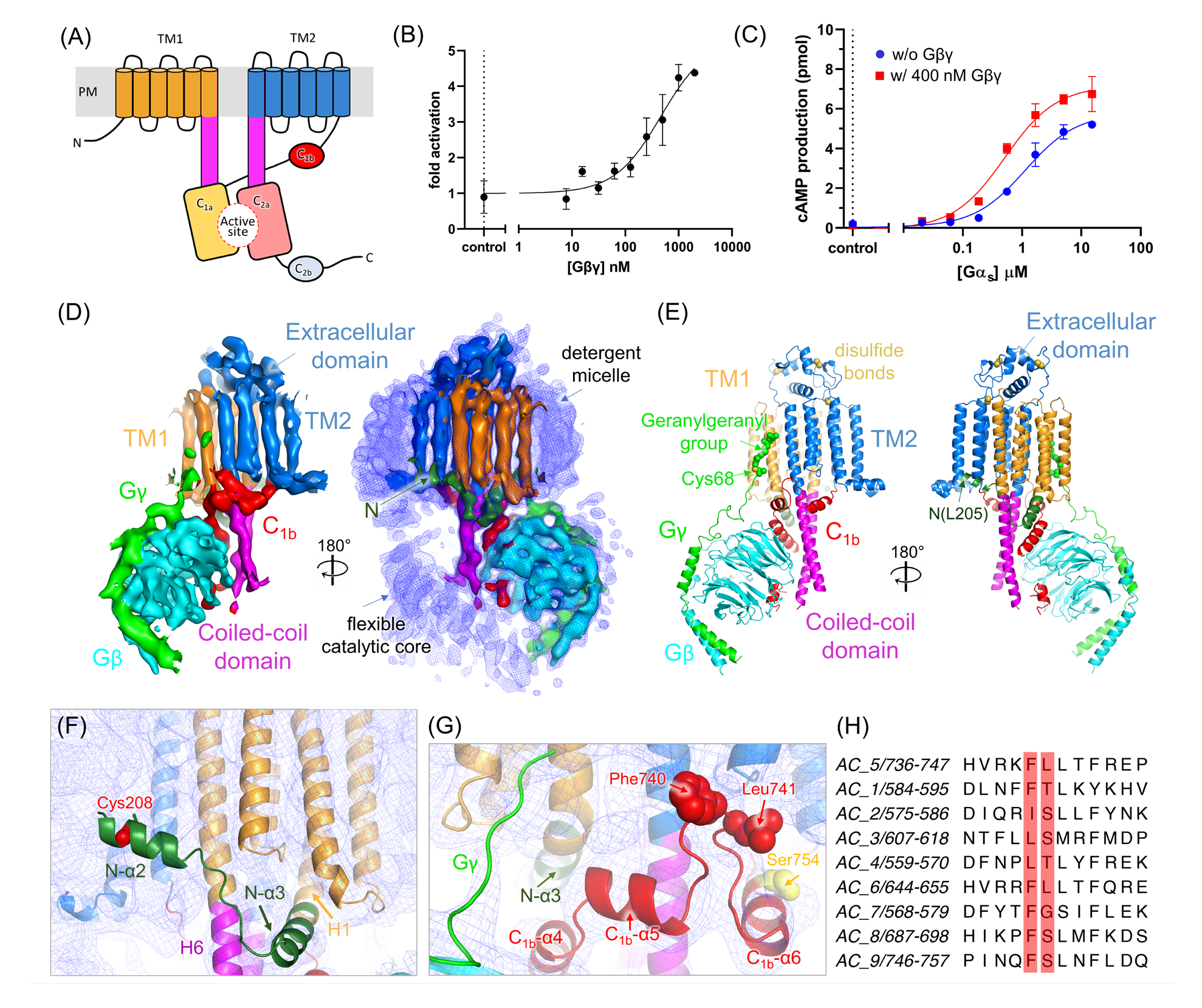
Characterization of AC5 and its complex with Gβγ. (A) Schematic of a typical mAC. TM: transmembrane domain; C_1a_, C_1b_, C_2a_, C_2b_: cytosolic domains; PM: plasma membrane. C2b is only 7 residues long in AC5. (B) Activation of AC5 by Gβγ. AC activity of purified AC5 was determined in the presence of 10 mM MgCl_2_, 0.5 mM ATP, and 100 nM Gα_s_·GTPγS at the indicated Gβγ concentrations. Data are the mean ± SD (n=3). (C) AC activity of purified AC5 was determined in the presence of 10 mM MgCl_2_, 0.5 mM ATP with (▪) or without (•) 400 nM Gβγ at the indicated Gα_s_ concentrations. Data shown are the mean ± SD (n=3). (D) Map and (E) corresponding atomic model for the AC5–Gβγ complex. Density for the catalytic core is however uninterpretable, indicating that it is not fixed relative to the coiled-coil domain. The O-methylcysteine and geranylgeranyl group at the C-terminus of Gγ, and predicted disulfide bonds in the extracellular domain are depicted as spheres, although there is no significant density for these modifications in the reconstruction. (F) Elements of the N-terminal domain. Density for the N-α3 helix and the linker between N-α2 and N-α3 is clearly visible. The N-α2 helix density merges with the micellar boundary, consistent with the presence of a high probability palmitoylation site at Cys208 (red sphere). (G) Region of C_1b_ wrapping around the micellar surface with the side chains of Phe740, Leu741, and Ser754 shown as spheres. Density is shown as blue mesh in panels F and G. (H) Sequence alignment of the linker between C_1b_-α5 and C_1b_-α6, with residues Phe740 and Leu741 in AC5 highlighted in red.

Advancements in cryo-electron microscopy (cryo-EM) have now enabled structure determination of large integral membrane protein complexes despite limited amounts of sample. In 2019, the cryo-EM structure of the full-length bovine AC9 was reported^8^, which revealed that the two integral membrane domains (TM_1_ and TM_2_) together form a 12- transmembrane helical bundle from which the C-terminal ends of two long helices (helix 6 and helix 12) protrude from the membrane to form a coiled-coil domain. This extended domain connects with the catalytic C_1a_ and C_2a_ domains and is analogous to coiled-coil domains observed in guanylyl cyclase (GC)^9^. However, the AC9 structure did not provide insights into the N-terminal or C_1b_ domains, which in other ACs are expected to be involved in binding A-kinase anchoring proteins^10^, Ca^2+^·calmodulin^5^, and Gβγ^6^. Furthermore, AC9 is only regulated by Gα_s_, rendering it unsuitable for the study of AC regulation by other signaling proteins^3^.

To explore isoform-specific regulation of mACs, we focused on human AC5 because its activity is regulated by various key signaling proteins in addition to Gα_s_, namely Gα_i/o_ and Gβγ^3^. Moreover, due to its significance in heart disease and dyskinesia, AC5 is an important therapeutic target. When overexpressed in transgenic mice, it results in the development of more severe cardiomyopathy in response to isoproterenol-induced oxidative stress relative to controls^11^. Furthermore, somatic gain-of-function mutations have been identified in *ADCY5* and are linked to familial dyskinesia^12, 13^. Understanding the unique regulatory mechanisms of AC5 could therefore facilitate the development of drugs with better selectivity and an understanding of human disease. In this study, our primary goal was to uncover mechanisms underlying the regulation of AC5 by Gβγ, a modulator of several mACs. Our structure of the AC5**–**Gβγ complex reveals large portions of the C_1b_ domain, and shows that Gβγ binds to both the coiled-coil and C_1b_ regions of the enzyme, suggesting that it could regulate the catalytic core allosterically by altering the twist of the coiled-coil, as is proposed in the regulation of GCs^14^. We also observed dimeric forms of AC5 under specific detergent conditions. Because Gβγ binding is incompatible with the homodimer interface, it suggests an additional regulatory mechanism wherein Gβγ releases Gα_s_-bound AC5 from a less active, homodimeric state.

## Results

### Validation of purified human AC5 and its regulation by Gβγ

Human AC5 was purified from mammalian cells and its functionality was confirmed using a cAMP production assay (**Fig. S1A**) and its ability to be activated by forskolin and Gα_s_·GTPγS, with EC_50_ values of 10 and 1 µM, respectively (**Fig. S1B and C**). Human Gβ_1_γ_2_ could also enhance AC5 activity after incubation with Gα_s_·GTPγS, as anticipated ^6, 15^ (**Fig. S2A**), with an EC_50_ value of 460 nM (**Fig. 1B, Table S1**). Pre-incubation of Gβγ with AC5 decreased the EC_50_ value for Gα_s_ 2-fold, accompanied by an increase in cAMP production (**Fig. 1C**), suggesting that Gβγ stabilizes AC5 in a configuration that optimizes the assembly of the C_1a_/C_2a_ catalytic domains in complex with Gα_s_. A direct AC5**–**Gβγ interaction was demonstrated via pull-down assay using anti-FLAG M2 affinity beads, and the size exclusion chromatography profile of the resultant complex was consistent with a homogenous 1:1 complex (**Fig. S2B and C**).

### Structure of the AC5–Gβγ complex

The purified AC5–Gβγ complex was incorporated into LMNG micelles and subjected to cryo-EM single particle analysis. The resulting 2D class averages revealed orthogonal views of the transmembrane (TM) domain, including a “top” view perpendicular to the membrane plane that clearly displays the 3×4 array of 12 transmembrane helices (**Fig. S3, Table S2**). A 7 Å EM reconstruction was generated from the selected particles which allowed a good fit with an AC5 model predicted by AlphaFold2^16^ and with prior crystal structures of Gβγ (**Fig. 1D and E**). The arrangement of the TM helixes in AC5 resembles those in the structure of AC9, with pseudo-2-fold symmetry and an extended coiled-coil domain formed by C-terminal extensions of helices 6 and 12 (H6 and H12) that reach ∼50 Å into the cytoplasm. It was not possible to model the C_1a_/C_2a_ catalytic domain, which is dynamic in this complex, although density is present (**Fig. 1D**, **E and Fig. S3**). Compared to AC9, AC5 has a more ordered extracellular domain formed primarily by its extended TM9-10 and TM11-12 loops (**Fig. 1D and E**). All cysteine residues within the extracellular domain of AC5 are configured such that they could form disulfide bonds (**Fig. 1E**), but only the Cys854-Cys904 disulfide might be conserved across mACs. The structure reveals additional elements of the N-terminal domain and C_1b_ that were not observed in AC9. The second predicted helical element in the AC5 N-terminus (N-α2, residues 205-217) is amphipathic and partially ordered at the micelle boundary, consistent with the presence of a high probability palmitoylation site at Cys208^17^ (**Fig. 1F**). N-α2 is connected by a ∼6 residue span to N-α3 (residues 224-237), which was observed in AC9 and packs at an angle against H6 of the coiled-coil domain, acting essentially as a bent N-terminal extension of the TM1 helix (**Fig. 1E and F**). Strong density wrapping around the coiled-coil domain like a belt near the micellar surface contains helices C_1b_-α4 (residues 712-726, which packs in parallel orientation against N- α3), C_1b_-α5 (residues 729-736), and C_1b_-α6 (residues 746-755, which harbors a PKA phosphorylation site at Ser754^18^) (**Fig. 1E and G**). This region was not observed in AC9, although predicted in AC9 by AlphaFold2^16^. An unusual 9 amino acid hairpin loop is found between C_1b_-α5 and C_1b_-α6 featuring Phe740 at its tip, whose side chain along with that of Leu741 is positioned to engage the hydrophobic phase of the micelle. This element is highly conserved among mACs (**Fig. 1G and H**).

### Engagement of Gβγ

The position of Gβγ is unambiguous in the reconstruction and buries ∼2700 Å^2^ of accessible surface area in the AC5–Gβγ complex (**Fig. 2A**). Density linking the C-terminus of G_γ_ with the surface of the detergent micelle is evident and was modeled as the geranylgeranylated C-terminus of the protein (**Fig. 1D**). A helical element in the C_1b_ region (modeled as C_1b_-α2, residues 690-696) packs against the central β- propeller domain of Gβγ (**Fig. 2A**), consistent with the top-scoring binding model predicted by AlphaFold Multimer^19^ (**Fig. S4A**). Mutation of residues in Gβ involved in the interface with C_1b_-α2 (Gβ-W99A, M101A, D186A, and N230A) were previously shown to reduce Gβγ-mediated AC5 activation^6^. Of these, Gβ-Trp99 forms hydrophobic contacts with AC5-Leu433, Leu436, and Met696. Gβ-Met101 forms hydrophobic interactions with AC5-Met696; and Gβ-Asn230 is in position to form a hydrogen bond with the side chain of AC5-Lys691 (**Fig. 2B**). Other Gβγ-interacting proteins, including Gα^20^, GRK2^21^, and Prex-1^22^, bind to an overlapping region (**Fig. S4B-D**). In addition, the side chains of AC5- Met1029 and Leu1032 in H12 of the coiled-coil domain pack against with Gβ-Leu55 (**Fig. 2C**). The side chain of AC5-Glu1025 is positioned to make a favorable electrostatic interaction with Gβ-Arg52, and multiple residues in C_1b_-α4, including Phe719, Arg722, and Ala726, interact with residues along the side of the Gβ propeller, including the side chains of Gβ-Thr50, Arg52, and Phe335. Residues involved in Gβγ binding in the coiled-coil and C_1b_-α4 helix are highly conserved in mACs except for AC9, which is insensitive to Gβγ regulation^3^ (**Fig. 2D**). Therefore, it is possible that these various AC5 regions constitute a common binding site for Gβγ that is shared among all Gβγ-responsive AC isoforms. An element within the N-terminus of AC5 (residues 66-137) has been reported to be involved in binding to Gβγ^6^, however this region is not evident in our structure (**Fig. 2A**).

**Figure 2.**
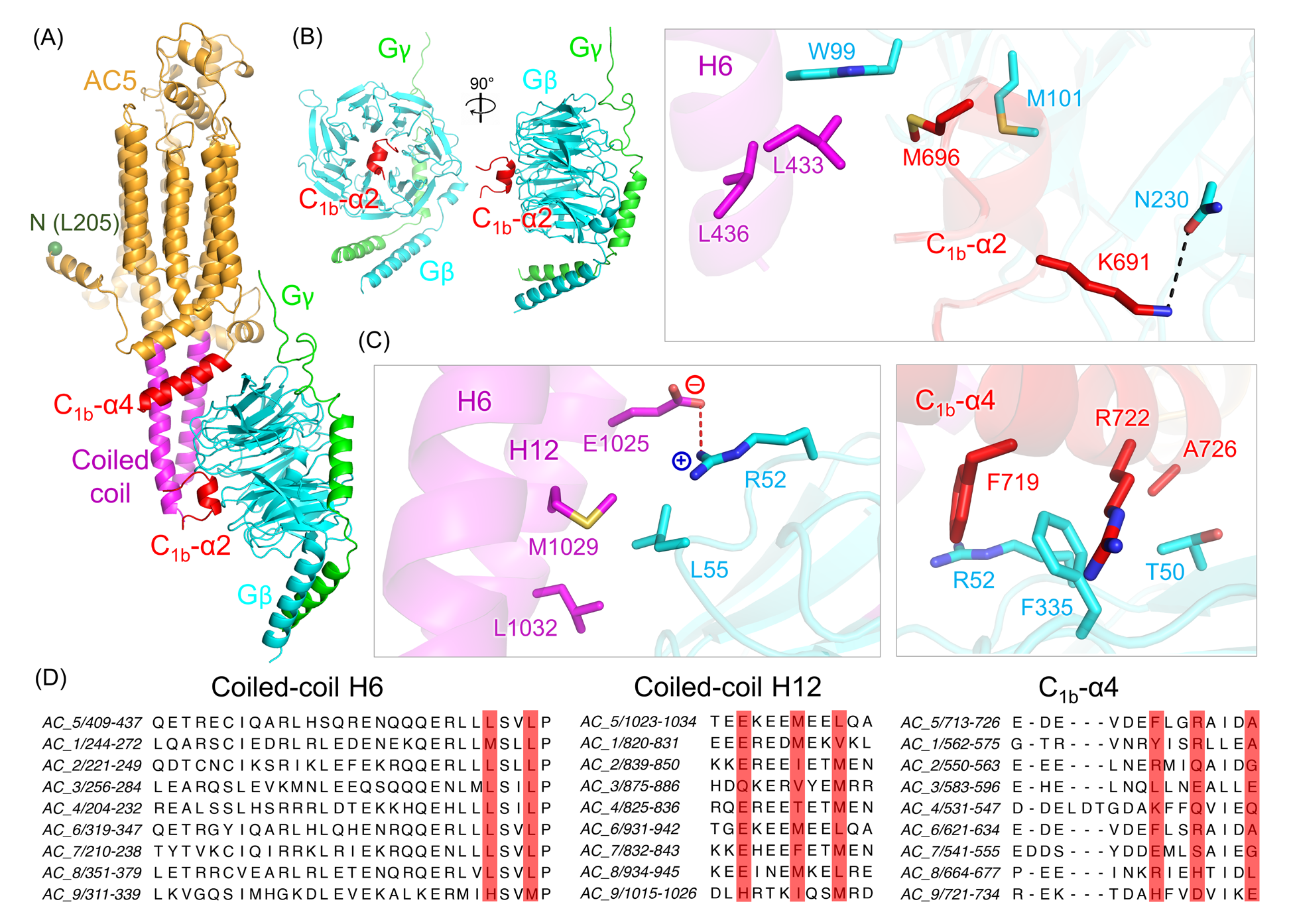
The AC5–Gβγ interface. (A) EM model of the AC5–Gβγ complex, with AC5 shown mostly in orange and the first modeled amino acid (residue 205) highlighted by a green sphere. The coiled-coil and C_1b_ domains, which are the primary regions of AC5 involved in Gβγ binding, are colored magenta and red, respectively. Gβ and Gγ are shown in cyan and green, respectively. (B) The binding of C_1b_-α2 in AC5 to the core of Gβ (left panel), and the potential interactions between H6 in the coiled-coil domain (magenta) and C_1b_-α2 (red) in AC5 with hotspot residues in Gβ (right panel). A black dashed line indicates a putative hydrogen bond. (C) The interaction of H12 in the coiled-coil domain (magenta, left panel) and C_1b_-α4 (red, right panel) in AC5 with Gβ. The residues involved in binding are represented in sticks and colored based on atom types. A red dashed line indicates a potential ionic interaction. (D) Sequence alignment of Gβγ interacting regions in the coiled-coil domain and C_1b_-α4 in human mACs. The residues involved in binding in AC5–Gβγ complex are highlighted with red boxes.

### Confirmation that the observed interface plays a role in Gβγ-mediated AC5 activation

First, the C_1b_-α2 region was deleted to create AC5_Δ687-708_ (**Fig. S5**) which exhibited significantly higher activity than the full-length protein under all tested conditions (**Fig. 3A**) and indicates that the C_1b_-α2 region in AC5 acts as an autoinhibitory element. Consistent with this result, deletion of residues Lys694-Met696 (Δ9bp) was previously shown to significantly increase AC5 response to Gα_s_ and forskolin-mediated stimulation in cell-based assays^23^. AC5_Δ687-708_ is relatively insensitive to Gβγ activation and exhibits at least a 6-fold reduction in EC_50_ (**Fig. 3B, Table S1**). In membranes from an HEK293 cell line lacking endogenous AC3 and AC6 (HEK-ACΔ3/6)^24^, we showed that overexpressed AC5 was activated by purified Gβγ with an EC_50_ value of 40 nM (**Fig.3C, Table S1**). In contrast, the EC_50_ value for AC5_Δ687-708_ was at least 100-fold higher and had lower maximal activity (**Fig.3C, Table S1**).

**Figure 3.**
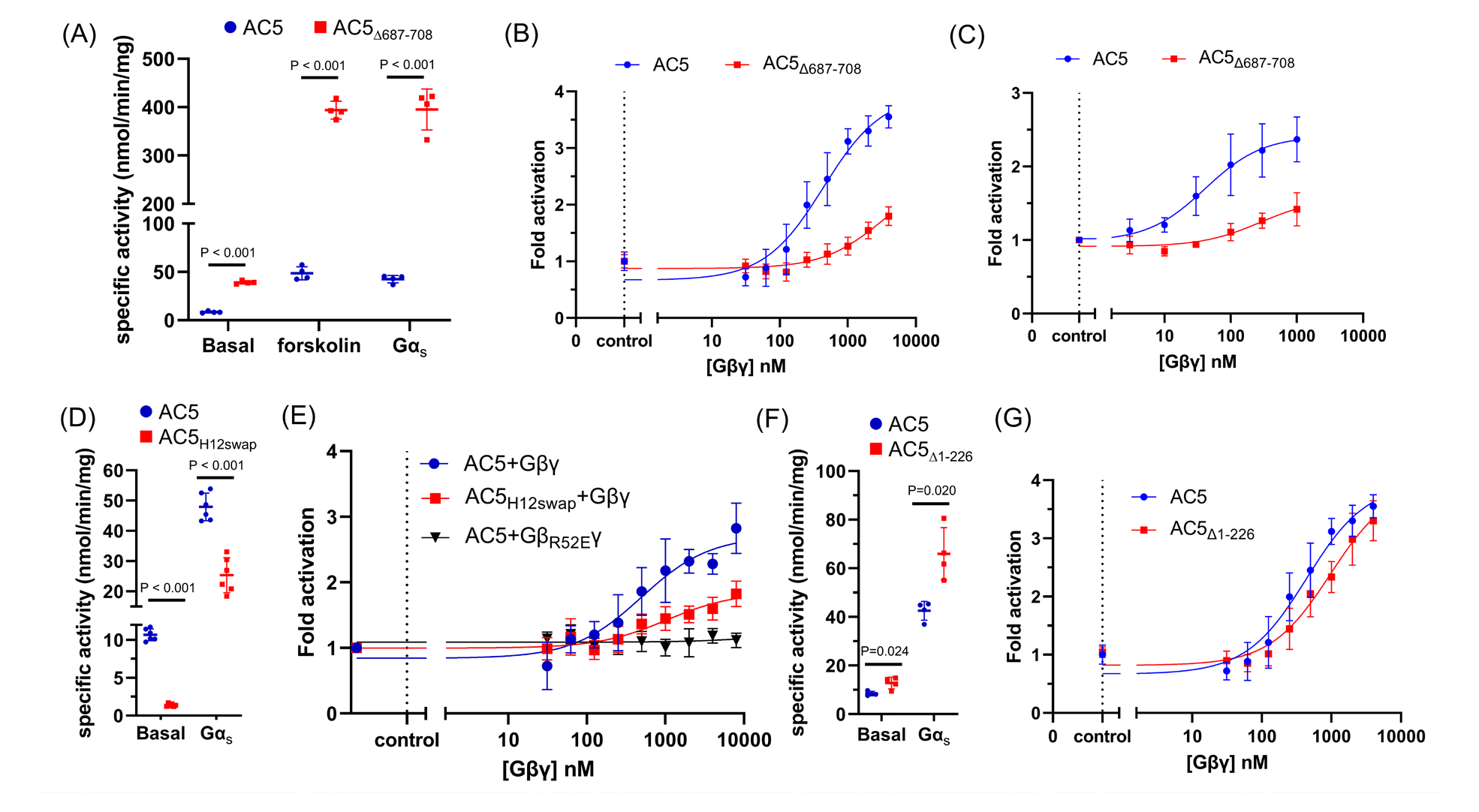
The C_1b_-α2 and H12_1023-1034_ regions in AC5 strongly contribute to Gβγ-mediated AC5 activation. (A) Specific activity of AC5 (•) and AC5_Δ687-708_ (▪) under basal conditions and after forskoloin or Gα_s_ stimulation. (B) Gβγ dose-response curves for AC5 (•) and AC5_Δ687-708_ (▪). (C) AC activity of AC5 and AC5_Δ687-708_ in HEK-ΔAC3/6 membranes in the presence of 10 mM MgCl_2_, 100 µM ATP, and 30 nM Gα_s_·GTPγS at the indicated Gβγ concentrations. Data presented are the mean ± SD (n=3). (D) Specific activity of AC5 (•) and AC5_H12swap_ (▪) under basal conditions and after Gα_s_ stimulation. (E) Gβγ dose-response curves for AC5+Gβγ (•) AC5_H12swap_+Gβγ (▪), and AC5+Gβ_R52E_γ (▾). (F) Specific activity of AC5 (•) and AC5_Δ1-226_ (▪) under basal conditions and after Gα_s_ stimulation. (G) Gβγ dose-response curve for AC5 (•) and AC5_Δ1-226_ (▪). In (A), (D) and (F), AC5 activity (mean ± SD) was measured in the presence of 10 mM MgCl_2_, 0.5 mM ATP with either 50 µM forskolin or 250 nM Gα_s_·GTPγS. In (B), (E) and (G), the Gβγ dose-response curves were determined in the presence of 10 mM MgCl_2_, 0.5 mM ATP, and 100 nM Gα_s_·GTPγS at the indicated Gβγ concentrations, with data presented as mean ± SD (n=3).

The binding interface between H12 in AC5 and Gβ-Arg52 (**Fig. 2C**) was also assessed. We replaced residues 1023-1034 of H12 in AC5 with those of AC9 to generate AC5_H12swap_ (**Fig. 2D and Fig. S5**). Compared to wild-type (WT), AC5_H12swap_ exhibited a significant reduction in basal and Gα_s_-stimulated activity (**Fig. 3D**). Dose-response curves also revealed diminished activation of AC5_H12swap_ by Gβγ (**Fig. 3E, Table S1**). The Gβ_R52E_γ variant (**Fig. S5**) exhibited more severe defects (**Fig. 3E**). To assess the effect of the AC5 N-terminus on Gβγ regulation, AC5_Δ1-226_ was purified (**Fig. S5**) and evaluated. The variant displayed similar activity compared to the full-length protein (**Fig. 3F**) and retained its susceptibility to Gβγ modulation with only a modest increase in the EC_50_ value (**Fig. 3G, Table S1**). This data aligns with previous evidence that the AC5 N-terminus contributes to binding but not activation by Gβγ^6^.

### Disease-related somatic mutations in AC5 cluster at the Gβγ binding interface

Multiple single nucleotide polymorphisms (SNPs) have been identified in human *ADCY5* and are linked to various disorders such as diabetes^25^, obesity^26^, and central nervous system disorders^27^ (**Fig. 4A**). Many of the SNPs are in regions known to be involved in modulation of AC activity, including six in the coiled-coil domain (**Fig. 4B**), six in C_1b_ (**Fig. 4C**), and ten in the catalytic domains (**Fig. 4D**). For example, the SNPs Y233H within N-α3, which packs against the coiled-coil domain, and D1015E within the coiled-coil domain are correlated to neurological disorders^28, 29^ and could affect the packing of the coiled-coil domain and consequently modulation of the catalytic core. Many SNPs involve residues located near the Gβγ binding interface (**Fig. 4A-D**). Among them, R418W, A726T, M1029K, and Δ9bp are associated with familial dyskinesia and represent gain-of-function mutations^12, 23^. We evaluated how Gβγ affected the AC activity of R418W, M1029K, and Δ9bp expressed in HEK-ACΔ3/6 cell membranes. All exhibited significant defects in Gβγ activation. R418W and Δ9bp had significantly reduced AC5 activation by Gβγ and dramatically higher EC_50_ values (**Fig. 4E, Table S1**). M1029K had similar maximal responsiveness but a right shift in the EC_50_ value (**Fig. 4E, Table S1**). Thus, gain-of-function in these variants strongly correlates with loss of Gβγ-induced activation, suggesting that the SNPs and Gβγ activate AC5 via the same mechanism.

**Figure 4.**
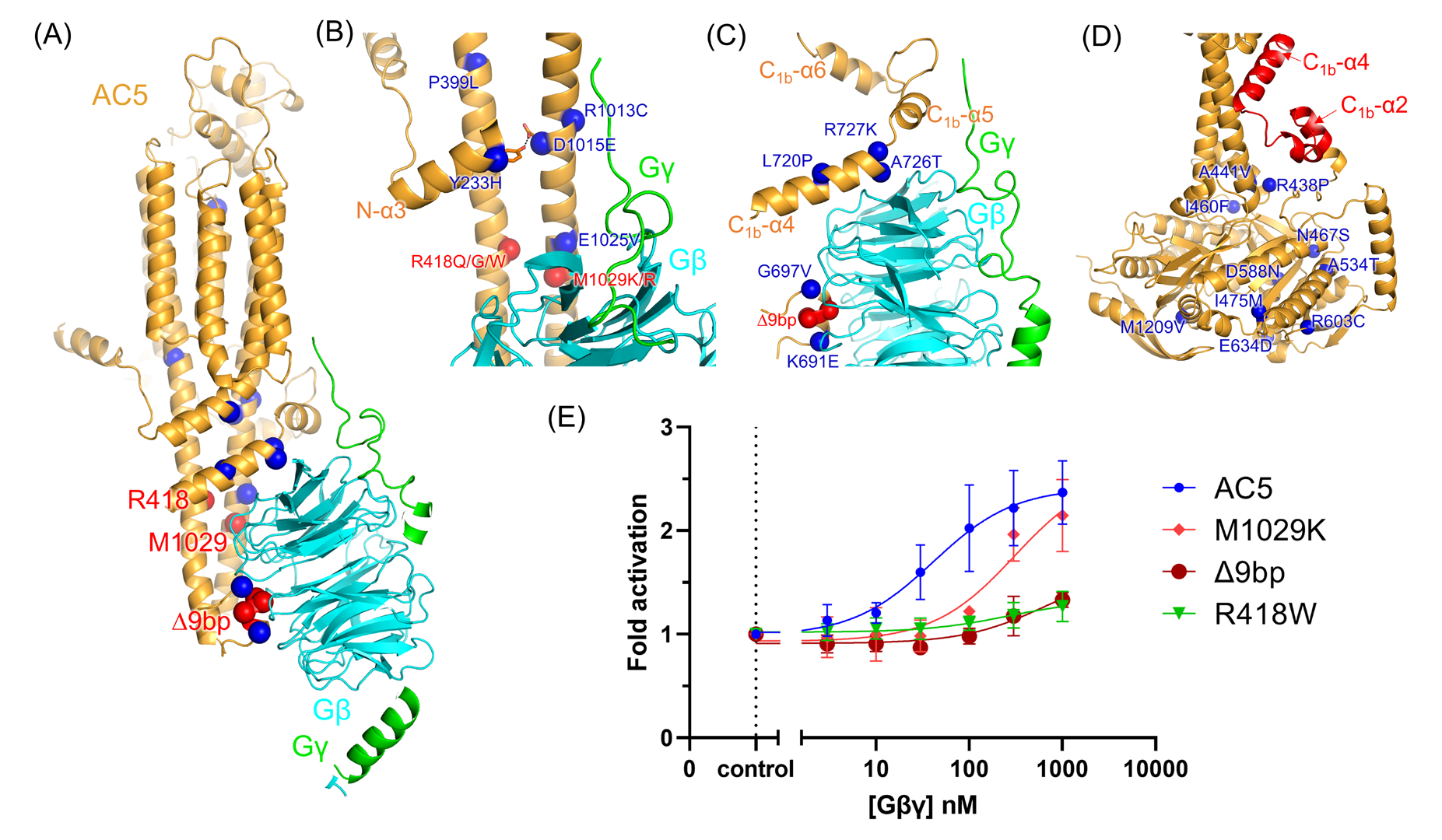
Location of disease associated AC5 mutations and impaired responses to Gβγ of those associated with familial dyskinesia. (A) Cartoon representation of the AC5-Gβγ complex. Somatic mutations identified in AC5 linked to disease are depicted as spheres at their Cα positions. AC5 SNPs that were examined in this study are shown in red. The somatic mutations located at (B) coiled-coil domain, (C) C_1b,_ and (D) catalytic core (from a AlphaFold2 model) are indicated with spheres. (E) Gβγ dose-response curves for AC5 variants in HEK-ΔAC3/6 cell membranes. AC assays were in the presence of 10 mM MgCl_2_, 100 µM ATP, and 30 nM Gα_s_·GTPγS at the indicated Gβγ concentrations. Data shown are mean ± SD (n=3).

### Dimers of AC5 form in the absence of bound Gβγ

We also sought to structurally characterize the Gα_s_–AC5·forskolin–Gβγ complex via cryo-EM analysis in LMNG micelles. The protein in these micrographs however displayed a severe degree of aggregation (**Fig. S6**). To address this concern, the Gα_s_–AC5·forskolin–Gβγ complex was instead reconstituted in GDN micelles. A dataset containing approximately 1,400 micrographs was collected, and the resulting 2D class averages revealed a heterogeneous population consisting of a monomeric AC5–Gβγ complex and, surprisingly, a dimeric AC5 complex with no obvious bound Gβγ (**Fig. S7**). To improve the map of the dimeric AC5, two-fold symmetry was imposed. Only one orientation of full-length AC5 (as predicted by AlphaFold2) is consistent with the observed 2D class averages, in that the orientation of the extracellular domains of each monomer and the side views of the 3×4 array of TM spans are distinct in orthogonal views (**Fig. 5A**). In this orientation, the hairpin loop between C_1b_-α5 and C_1b_-α6 forms the primary interaction surface between the TM domains (**Fig. 5A and B**), burying 2400 Å^2^ of accessible surface area. The resulting model is incompatible with Gβγ binding, explaining why there might be two classes of molecules in our data set. Keeping the inter-subunit distance consistent with the spacing of the coiled-coil domains evident in the 2D averages (**Fig. 5A**) would lead to overlap of the AlphaFold2 modeled catalytic core C_1a_ and C_2a_ domains, which likely explains the poor density in this region. Slight alteration of the two catalytic cores permits a reasonable fit to the density with contacts that seem to be mediated by the extended α1-α2 loops of the C_2a_ domains (**Fig. 5A and C**). Given that the α1-α2 loop is an element that also engages Gα_s_, it suggests that the homodimer is also not fully compatible with Gα_s_ binding. However, the 3D reconstruction and 2D class averages reveal blurred density consistent with the presence of Gα_s_ near this element (**Fig. 5A and Fig. S7**), which we interpret to mean that there are multiple configurations of the catalytic domains in our dimeric ensemble, some that are bound to Gα_s_ and some that are not. Consistent with this, observation of AC5 homodimers is dependent on detergent choice (**Fig. S6**). For example, the AC5–Gβγ–Gα_s_ complex isolated in LMNG yielded particles similar to the AC5–Gβγ complex (including its dynamic catalytic core) and no homodimers (**Fig. S6**). GDN is a glycosylated analog of cholesterol, a lipid that is abundant in biological membranes, and thus we conjecture that the observed AC5 homodimers are stabilized by cholesterol and that they could be biologically relevant. Indeed, there is strong evidence that the closely related isoform AC6 can be regulated by cholesterol^30^, and that AC5/6 are localized to cholesterol rich domains^31^.

**Figure 5.**
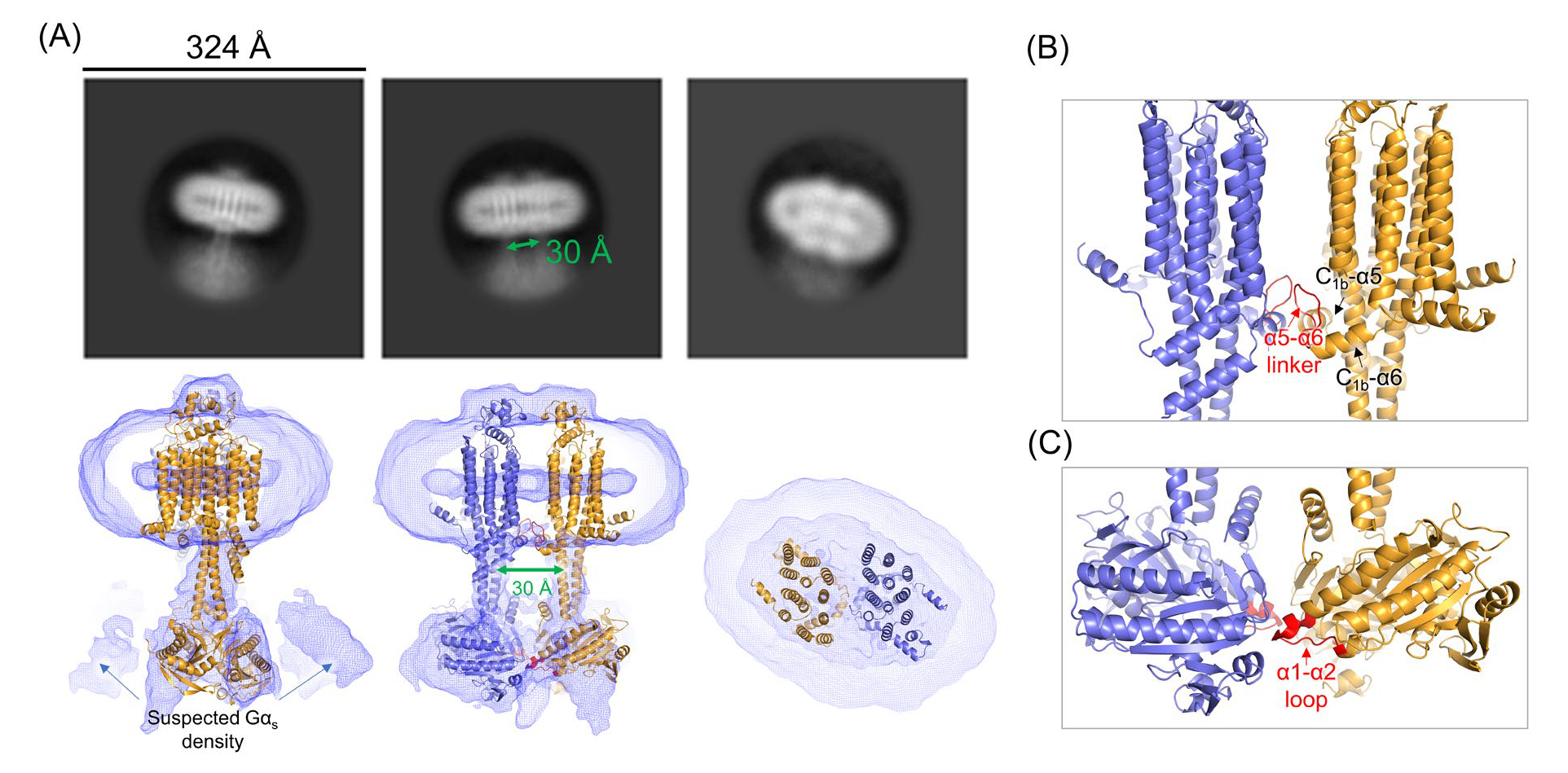
Homodimerization of AC5 could be linked to regulation by Gβγ. (A) Representative 2D class averages (top panel, box size = 324 Å, mask = 180 Å) and the corresponding views of the 3D reconstructed model (bottom panel). The intermonomer distance between the coiled-coil domain is indicated by double arrows. AC5 is depicted in cartoon representation, and the regions involved in the dimer interface are highlighted in red in the bottom middle panel. The dimerization interface seems mediated by the α5-α6 linker in C_1b_ and the α1-α2 loop in C_2a_, as shown in (B) and (C), respectively.

## Discussion

The molecular mechanisms underlying isoform-specific regulation of individual mACs have remained elusive for several decades. Gβγ modulates the activity of all mAC isoforms except AC9 in either a stimulatory (AC2, 4, 5, 6, and 7) or inhibitory (AC1, 3, and 8) manner^3^. Among them, AC2 has been the most extensively studied because of its robust conditional activation in response to Gβγ. Through domain swapping, mutagenesis, and peptide competition studies, the binding interface between AC2 and Gβγ has been variously reported to involve sites such as the “PFAHL motif” in C_1b_^32, 33^, the QEHA region in C_2a_^34^, the KF loop in C_2a_ (residues 926-934)^35^, and two additional sites in C_1_ (C_1a_ residues 339-360 and C_1b_ residues 578-602)^36^. However, none of the sites located in the C_1a_ or C_2a_ domains are consistent with the Gβγ binding site we observe in AC5, which instead involves the coiled-coil domain and helical elements within C_1b_. In fact, the C_2a_ domain and its “QEHA” motif is quite distant from the observed interface as modeled. However, we note that our best data was derived from samples in the absence of an AC activator, and it is possible that additional binding sites, such as those located in the N-terminus or catalytic C_1a_ and C_2a_ domains of AC5 would become evident when the enzyme is fully activated. However, given the surface of Gβγ that is engaged in our complex (**Fig. 1D and 2A**), there is little room left for additional canonical interactions^37^. Even so, it remains possible that the N-terminus (or other C1 regions) stabilize interactions of C_1b_ with Gβγ indirectly.

The low resolution of our reconstructions rendered interpretation of the AC5 density adjacent to Gβ-Trp99 somewhat ambiguous, as it could correspond to several different regions in C_1b_. The modeling we report is however supported by predictions from AlphaFold Multimer^19^ and the fact that mutations of residues in the 687-708 region reduce or eliminate Gβγ activation (**Fig. 3B and C**). The AC5 interface with Gβγ we model also harbors disease-associated mutations in AC5, including R418Q/W/G, K691E, R695E, G697V, Δ9bp, L720P, A726T, R727K, E1025V, and M1029K/R^3^. We show here that at least some of these mutations exhibit loss of Gβγ activation (**Fig. 4E**). We note that the PFAHL motif in C_1b_ proposed to be the Gβγ binding motif in AC2^32, 33^ is not well conserved in either AC5 or its close homolog AC6 (**Fig. S8**). We therefore speculate that Gβγ binds similarly to all mACs because the surface topologies revealed here will be similar among all isoforms and the position of Gβγ is also constrained by its distance to the membrane surface, as dictated by the prenylated C-terminus of Gγ. However, the specific elements that bind to the canonical hotspot^37^ of Gβγ may be distinct among the Gβγ-regulated isoforms, which may in turn lead to either activation or inhibition. Indeed, our cryo-EM model of the AC5-Gβγ complex seems somewhat consistent with a proposed docking model of AC8-Gβγ (wherein Gβγ would be inhibitory) suggested by crosslinking experiments^38^.

Many, if not all, of the mutations involved in the Gβγ interface in AC5 are gain-of-function, indicating that they somehow relieve autoinhibition of the enzyme. They also reduce or eliminate activation by Gβγ, implying that the mutations and the binding of Gβγ are involved in a similar mechanism of activation. Although our data is not yet sufficient to reveal details about what this autoinhibited state looks like, they point to a number of potential mechanisms. The first is that Gβγ may drive AC5 into an exclusive monomeric state that has higher specific activity than the homodimer (**Fig. 6**), perhaps because Gα_s_ site is partially occluded, or the catalytic core is not optimally aligned within the homodimer. Because the gain-of-function mutations pack near or are in the coiled-coil domain, it is possible they may achieve the same result via allostery. Such could explain the dependence of Gβγ activation on the presence of Gα_s_ or forskolin because the catalytic core remains misconfigured in either homodimer or monomer form without them. The second mechanism is that the α2 region of C_1b_, and/or surrounding elements, engages these gain-of-function regions to render AC5 less active. Gβγ-binding sequesters these autoinhibitory elements and thereby enhances activity, so long as Gα_s_ or forskolin are also present (**Fig. 6**).

**Figure 6.**
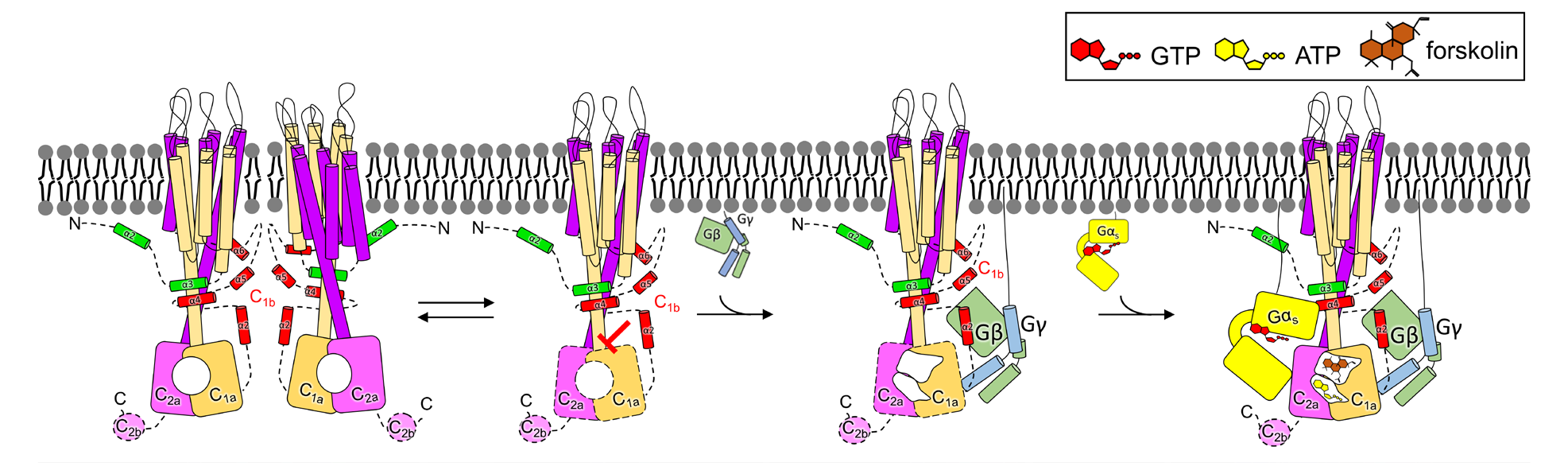
Proposed model for Gβγ-mediated AC5 activation. Our study suggests that AC5 can exist in a monomer-dimer equlibrilum, with Gβγ exhibiting a preference for the monomer. Our biochemical data further indicates that the C_1b_ domain contains an autoinhibitory element. We propose that Gβγ binding to C_1b_, AC5 is released from its autoinhibitory state, which may correspond to the homodimer, making it ready for catalysis. Domains outlined with dashes indicate that they are poorly ordered or unstructured in currently available structures.

The ability of AC5 to form a homodimer was demonstrated in COS-7 cells using BiFC^2^, and mass photometry analysis of our purified AC5 protein in DDM micelles also evidenced a mixed population of monomer, dimer, and higher order oligomers (data not shown). The high conservation of the hairpin loop between C_1b_-α5 and C_1b_-α6 (**Fig. 1H**) may indicate that it mediates homo or heterodimerization of other isoforms, although there is as of yet no structural evidence for homodimerization of AC8 even though their structures were determined in the presence of GDN^38^. Dimerization of mACs has previously been proposed as a regulatory mechanism^39^. The existence of dimeric mAC was initially suggested by hydrodynamic analysis of detergent solubilized calmodulin-sensitive AC isolated from bovine cerebral cortex^40^. Further evidence was obtained through the pull-down of an active AC complex using an antibody recognizing the C-tail of AC1 from Sf9 cells co-expressing an inactive mutant and C-terminally truncated AC1^41^. The existence of homodimeric AC8 was reported based on several experiments. In cell-based assays, it was detected through co-immunoprecipitation, FRET, and functional assays^42^. It was also observed in purified protein from size exclusion chromatography and crosslinking experiments^38^. A functional heterodimeric AC2/5 complex was demonstrated in HEK293 cells and whole mouse heart extracts by co-immunoprecipitation, confirming their presence physiologically^43^.

A recent preprint described the structures of AC8–Gα_s_^38^ in both GDN micelles and lipid nanodisc. Comparison of the structures of AC5, AC8, and AC9 reveal structural features that are both shared and unique. The N-terminal domain of mACs have varying lengths (**Fig. S9A**) and are disordered in all three reported structures, except for N-α3 (**Fig. 1D and Fig. S9B**). Our structure of AC5 shows a further N-terminal extension (**Fig. 1D and E**). The extracellular domain of AC5, like that of AC8, is predicted to have a pocket consisting of hydrophobic residues and a negatively charged surface, but the size of the binding pocket in AC5 is much smaller^44^ (**Fig. S9C**). The 12 TM helixes of AC5 are arranged similarly to AC9, but orientation of the coiled-coil domain in AC5 is somewhat different (**Fig. S9D**). We note that the structures of AC8 and AC9 were determined when they were fully activated, while our AC5–Gβγ complex was determined without any activators. However, the catalytic core of AC5 seems highly mobile relative to the TM spanning and coiled-coil domains regardless of its activation status.

In summary, our cryo-EM structure of the AC5–Gβγ complex revealed that non-conserved elements in the coiled-coil and C_1b_ domains of AC5 form the two major binding interfaces for Gβγ. The primary sequence of the C_1b_ domain is poorly conserved across isoforms (**Fig. S8**), consistent with its crucial role in isoform-specific regulation, and our data strongly suggests that it functions as an auto-inhibitory element that could either be displaced by activators or reinforced by inhibitors among the various mACs. Future studies of other regulatory complexes of AC5, such as with Gα_i_, will undoubtedly reveal additional regulatory complexity, but also unique features that might be exploited for isoform specific inhibition.

## Material and methods

### Reagents

Chemicals utilized in this study include forskolin from AvaChem Scientific, GDP and GTPγS from Sigma-Aldrich, n-dodecyl-β-D-maltoside (DDM) from Glycon, cholesteryl hemisuccinate (CHS) from Sigma-Aldrich, and lauryl maltose neopentyl glycol (LMNG), glyco-diosgenin (GDN), and CHAPS from Anatrace.

### Cloning of AC5 variants

All AC5 variants are generated by PCR using full-length human AC5 (hAC5, NCBI reference sequence: NM_183357.2) as a template and primers listed in **Table S3**. The resulting PCR products were then digested with XhoI and KpnI and cloned into a BacMam expression vector (pEZT-BM, Addgene plasmid # 74099)^45^. To facilitate protein purification, all constructs were designed with His_10_ and FLAG tags at the N-terminus and a 1D4 tag at the C-terminus. Expression vectors for disease-related gain of-function mutations were obtained from a previous study^23^.

### Expression and purification of human AC5

hAC5 was expressed in Expi293F GnTI^-^ (ThermoFisher) cells infected with baculovirus carrying hAC5 expression variants. The baculovirus was produced using the Bac-to-Bac Baculovirus expression system (Invitrogen) in *Spodoptera frugiperda* (Sf9) cells. To generate the bacmid, pEZT-BM encoded hAC5 was transformed into DH10Bac cells. The extracted bacmid was transfected into suspension Sf9 cells using FuGene to generate a P0 virus. The supernatant containing P0 virus was harvested five days post-transfection by pelleting down the cells with centrifugation at 3,000 g for 5 mins. P0 virus was further amplified twice in suspension Sf9 cells (infected at 2×10^6^/ml) to generate a P2 virus. The P2 virus was separated from the cell pellet by centrifugation at 3,000xg for 20 mins, filtered through 0.22 µM filters and used for mammalian cell infection. Suspension Expi293F GnTI^-^ cells were cultured at 37°C with 7% CO_2_ and shaking at 120 rpm. Once the cell density reached 2.5-3×10^6^/ml, the P2 virus was added into the cells with a multiplicity of infection of 1. After 18 hours of infection, 5 mM valproic acid was added into the culture, and the cells were returned to the incubator for an additional 48 hours before harvesting. The cell pellets were collected by centrifugation at 3,000xg for 20 minutes and stored at −80°C.

To lyse the cells, cell pellets were resuspended in PBS buffer supplemented with protease inhibitors (10 μg/ml phenylmethylsulfonyl fluoride and 1 μg/ml leupeptin). The cells were then homogenized using a Dounce homogenizer and lysed by passing them twice through an Emulsiflex (500-750 psi). The membrane fraction was collected by ultracentrifugation at 186,000xg (Ti45 rotor) for 45 minutes. The collected membrane was then re-suspended with PBS supplemented with 200 mM NaCl and 20 % glycerol. AC5 was extracted from the membrane by incubating with 1% DDM and 0.1 % CHS at 4°C for 3 hours on a stir plate. The insoluble fractions were removed by ultracentrifugation at 186,000xg for 45 minutes. The resulting supernatant was incubated overnight with anti-FLAG M2 magnetic beads (Sigma-Aldrich) at 4°C. The magnetic beads were then collected with a magnetic rack separator and washed three times with 20 bed volumes of PBS supplemented with either 0.025 % LMNG or 0.05 % GDN. AC5 was eluted three times by incubating with one column volume of elution buffer (wash buffer supplemented with 0.1 mg/ml 3x FLAG peptide) at 4°C for 20 minutes. For the AC5–Gβγ and AC5–Gβγ– Gα_s_ complexes, excess amounts of Gβγ and Gα_s_·GTPγS protein were added to the AC5- bound anti-FLAG M2 magnetic beads after the initial wash, and the mixture was incubated at 4°C for 2 hours. The magnetic beads were washed again to remove unbound Gβγ and Gα_s_ with wash buffer, and the complex was eluted with elution buffer. The eluted complexes were then concentrated to ∼0.5 mg/ml (as judged by SDS-PAGE) for cryo-EM analysis.

### Expression and purification of Gβγ

Geranylgeranylated human Gβ_1_γ_2_ protein with a N-terminal His_6_ tag on Gβ was purified from Sf9 cells as described previously^22^. Briefly, the cells were harvested two days after infection and lysed with an Emulsiflex C3. The membrane fraction was collected by ultracentrifugation (186,000 g for 40 mins) and solubilized with 1 % sodium cholate in buffer A (20 mM HEPES pH 8, 100 mM NaCl and 1 mM MgCl_2_). The insoluble fraction was removed by ultracentrifugation, and the supernatant was loaded onto Ni^2+^ resins (HisPur^TM^ Ni-NTA resin, ThermoFisher) equilibrated with buffer A. After washing with 20 column volumes of buffer B (20 mM HEPES pH 8, 100 mM NaCl, 1 mM MgCl_2_, 10 mM CHAPS and 10 mM imidazole pH 8), the protein was eluted with 5 column volumes of buffer C (20 mM HEPES pH 8, 100 mM NaCl, 1 mM MgCl_2_, 10 mM CHAPS, and 150 mM imidazole pH 8). The eluted protein was dialyzed against buffer D (20 mM HEPES pH 8, 0.5 mM EDTA, 2 mM MgCl_2_, 10 mM CHAPS, and 1 mM DTT) and loaded onto an anion exchange column (Q column) equilibrated with buffer D. The column was washed with buffer D until the UV_280_ signals were stable. Gβγ protein was eluted with a linear gradient to 30 % buffer E (20 mM HEPES pH 8, 1 M NaCl, 0.5 mM EDTA, 2 mM MgCl_2_, 10 mM CHAPS, and 1 mM DTT). The fractions were collected and pooled based on their purity on SDS-PAGE. The pooled fractions were then concentrated and further purified by size exclusion chromatography (SEC) using buffer F (20 mM HEPES pH 8, 100 mM NaCl, 0.5 mM EDTA, 2 mM MgCl_2_, 10 mM CHAPS, and 1 mM DTT). Peak fractions from SEC were pooled, fresh-frozen in liquid nitrogen, and stored at −80°C until use.

### Expression and purification of Gα_s_

A published protocol^46^ was followed for the expression and purification of Gα_s_ in *E. coli*. Briefly, the pQE-6 vector encoding His-tagged Gα_s_ was co-transformed with pREP4 vector into the BL21/DE3 strain. A single colony was used to inoculate 50 ml Luria-Bertani broth supplemented with 100 μg/ml carbenicillin and 50 μg/ml kanamycin and grown overnight at 37°C. The following day, 5 ml of overnight culture was added into 500 ml terrific broth supplemented with 100 μg/ml carbenicillin and 50 μg/ml kanamycin and grown at 37°C until the optical density (O.D._600_) reached 0.5. Then, 30 μM IPTG and 1 μg/ml chloramphenicol were added into the culture to induce protein expression. The culture was grown at 30°C for 18 hours before harvest. The collected pellet was stored at −80°C until the day of purification.

To purify the protein, cells were resuspended in lysis buffer (50 mM HEPES pH 8, 1 mM EDTA, 2 mM DTT, 10 μg/ml PMSF, and 1 μg/ml leupeptin) and lysed with Emulsiflex C3. Cell debris was removed by centrifugation at 100,000xg for 40 minutes, and the supernatant was collected and loaded onto Ni resin (HisPur^TM^ Ni-NTA resin, ThermoFisher) equilibrated with lysis buffer. After loading the sample, the Ni resin was washed with 20 column volumes of buffer containing 50 mM HEPES pH 8, 500 mM NaCl, and 10 mM imidazole pH 8. The protein was then eluted with a buffer containing 50 mM HEPES pH 8, 100 mM NaCl and 300 mM imidazole pH 8. The eluted protein was analyzed with SDS-PAGE before dialyzing it against a buffer containing 50 mM HEPES pH 8 and 2 mM DTT at 4°C overnight. The dialyzed protein was loaded onto a Q column equilibrated with the dialysis buffer. The Q column was washed with the same buffer supplemented with 2 μM GDP until UV_280_ stabilized. The Gα_s_ protein was eluted with a linear gradient from 0 to 100 % of a buffer containing 50 mM HEPES pH8, 2 mM DTT, 2 μM GDP, and 500 mM NaCl. The fractions were collected and pooled based on purity by SDS-PAGE. The pooled fractions were concentrated and further polished with SEC in a buffer containing 50 mM HEPES pH 8, 10 mM MgSO_4_, 1 mM EDTA, 2 mM DTT, and 2 μM GDP. The peak fractions from SEC were collected, concentrated, and stored at −80 °C until use. Prior to assay, the purified Gα_s_ was incubated with an excess amount of GTPγS at 4 °C overnight for nucleotide exchange.

### AC assays

To determine AC activity of purified AC5 variants, 20 nM AC5 was mixed with an equal volume of assay buffer (20 mM MgCl_2_, 1mM ATP with either 20 μM forskolin or 200 nM Gα_s_·GTPγS, and 0.025 % GDN in PBS). The mixture was then incubated at room temperature for 15 minutes and subsequently inactivated by heating at 95 °C for 5 minutes. The amount of cAMP produced in each reaction was determined using the cAMP Gs dynamic kit (Cisbio), following the manufacturer’s instructions. Briefly, the heat-inactivated reaction mixture was diluted with PBS to ensure that the amount of cAMP remained within the range of the standard curve. The diluted reaction mixture (or standard) was then mixed with d_2_-labeled cAMP and anti-cAMP cryptate-labeled antibody provided in the kit in a 384-well plate (Corning, REF3824). The plate was sealed and incubated at room temperature for 1 hour, then measured at 620 nm (donor emission signal) and 665 nm (acceptor emission signal) using a homogeneous time resolved fluorescence (HTRF) compatible plate reader (FlexStation 3). The HTRF ratio of the acceptor and donor emission signals was calculated for each well. To generate a standard curve, HTRF ratios were plotted against known cAMP concentrations. The amount of cAMP produced in each sample was calculated by interpolation from the standard curve. To determine the EC_50_ of different regulators (Gα_s_·GTPγS, forskolin, and Gβγ), serial dilutions of each regulator were prepared and assayed as described above. For measuring Gβγ regulation, AC5 was first incubated with Gβγ on ice for 20 minutes before adding the assay buffer.

### HEK-ΔAC3/6 cell transfections.

HEK-ΔAC3/6 cells^24^ were maintained in Dulbecco’s modified Eagle’s medium supplemented with 5% bovine calf serum, 5% fetal bovine serum, and 1% penicillin-streptomycin at 37°C with 5% CO_2_. Cells were transfected with the appropriate plasmids using Lipofectamine 2000 and used to prepare cell membranes 48 hours after transfection according to a previous established protocol^47^.

### Membrane AC assays

Membranes from HEK-ΔAC3/6 cells transfected with AC5 or indicated mutants were prepared and assayed as described^48^. Briefly, freshly prepared membranes were diluted in HMED (20 mM HEPES, pH 7.4, 2 mM MgCl_2_, 1 mM EDTA, and 1 mM DTT) and preincubated with the indicated concentrations of Gβγ for 5 min at RT. Gα_s_·GTPγS was added on ice and the assay was initiated with AC mix containing a pyruvate kinase regenerating system, 10 mM MgCl_2_, and 100 µM [^32^P-α]ATP. After 30 min at RT, reactions were terminated with a mix of 2.5 % SDS, 50 mM ATP, and 1.75 mM cAMP. Each reaction was subjected to column chromatography with [^3^H]cAMP added to monitor column recovery rates.

### Cryo-EM sample preparation and data collection

To make cryo-EM grids, 3.3 μl of purified AC5-Gβγ complex at 0.5 mg/ml was applied onto glow-discharged UltrAuFoil R1.2/1.3 300-mesh grids using EasiGlow at 15 mA for 90 seconds. The grids were then blotted for 3.5 seconds with filter paper and rapidly plunge-frozen in liquid ethane using a Vitrobot MK IV (Thermo Fisher Scientific). Micrographs of the grids were collected on a Titan Krios G1 electron microscope (FEI) equipped with a K3 summit direct electron detector (Getan) in the Purdue Life Science Cryo-EM facility. A dataset containing ∼1,700 images was collected in super-resolution mode with a pixel size of 0.54 Å, at a defocus range of 1 to 3 μm using Leginon. For each movie stack, 40 frames were recorded at a frame rate of 78 ms per frame and a total dose of 53.8 electrons/Å^2^.

### Cryo-EM data processing and model building

The image processing flowchart is shown in **Fig. S3**. Beam-induced motion was corrected, and the micrographs were binned twofold using MotionCor2 in Appion. The motion-corrected micrographs were then imported to CryoSPARC^49^ for estimating the contrast transfer function parameters using the CTFFIND4 module. A small set of particles were picked using blob picker to generate class averages, which were then used as templates for autopicking using template picker. Following several rounds of 2D classification, an initial model was generated using ab initio reconstruction, and the resulting 3D model was used for homogeneous refinement and nonuniform refinement. Local refinement was then carried out using a mask generated in RELION-3^50^ covering the transmembrane and helical domains of hAC5, as well as the entire molecule of Gβγ, to further improve map quality.

An hAC5 model obtained from the AlphaFold2 database and the crystal structure of Gβγ (PDB entry 6U7C) were rigid-body docked into the cryo-EM map using Chimera^51^. The resulting model was further fitted into the map using molecular dynamic flexible fitting (MDFF)^52^. The MDFF configuration files were generated using VMD^53^. During MDFF simulation, Gβγ was set as rigid with domain restraints. The MDFF simulation was conducted with a grid scaling value of 0.5 for 100 ps, followed by 3000 steps of energy minimization until convergence of the protein RMSD. The resulting model from MDFF was inspected, adjusted manually in COOT^54^, and refined using phenix.real_space_refine implemented in Phenix. To build the dimeric form of AC5, the AC5 model in the AC5–Gβγ complex was first fitted into the map based on the orientation of the transmembrane helices observed in the 2D class averages. The catalytic core model obtained from the AlphaFold database was then manually docked into the map and adjusted to alleviate steric overlap with the other monomer. The resulting dimeric AC5 model was then refined using Phenix.

### Statistics

Assays were conducted with at least three technical replicates (n=3). The significance of any differences observed in AC activity was determined using multiple unpaired t-tests implemented in GraphPad Prism 9. The dose response curves were fitted with a three-parameter logistic nonlinear regression model.

### Data availability

All data needed to evaluate the conclusions in the paper are present in the paper and/or the Supplementary Materials. The structures of the AC5–Gβγ complex and the AC5 homodimer have been deposited into the Protein Data Bank under accession codes 8SL3 and 8SL4, and the Electron Microscopy Data Bank under accession codes EMDB-40572 and 40573, respectively.

## Acknowledgements

The authors gratefully acknowledge the support of Purdue Life Sciences Electron Microscopy Facility from the Institute for Cancer Research, NIH grant P30 CA023168 and from the Indiana Clinical and Translational Sciences Institute (CTSI) FY21 CTSI PostDoc Challenge.

## Funding

This work was supported by NIH grants CA254402 (J.J.G.T), CA221289 (J.J.G.T.), CA023168 (J.J.G.T.), HL071818 (J.J.G.T.), GM60419 (C.W.D), GM145921 (C.W.D), DA051876 (V.J.W.), NS119917 (V.J.W.), and NS111070 (V.J.W.), the Walther Cancer Foundation (J.J.G.T.), and American Heart Association Postdoctoral fellowship 903842 (Y.C.Y.).

## Author contribution

Conceptualization: J.J.G.T., V.J.W., C.V.D. Y.C.Y.

Methodology: V.J.W., C.V.D. Y.C.Y., J.J.G.T.

Investigation: Y.C.Y., Y.L., C.L.C., and T.K.

Funding acquisition: J.J.G.T., V.J.W., C.V.D.

Project administration: J.J.G.T., V.J.W., C.V.D.

Supervision: J.J.G.T., V.J.W., C.V.D.

Writing—original draft: Y.C.Y.

Writing—review and editing: J.J.G.T., V.J.W., C.V.D., Y.C.Y., Y.L., C.L.C., and T.K.

## Competing interests

The authors declare that they have no competing interests.

## Material & correspondence

Requests for the plasmids and other reagents should be submitted to the corresponding author J.J.G.T.

---

**Figure S1.**
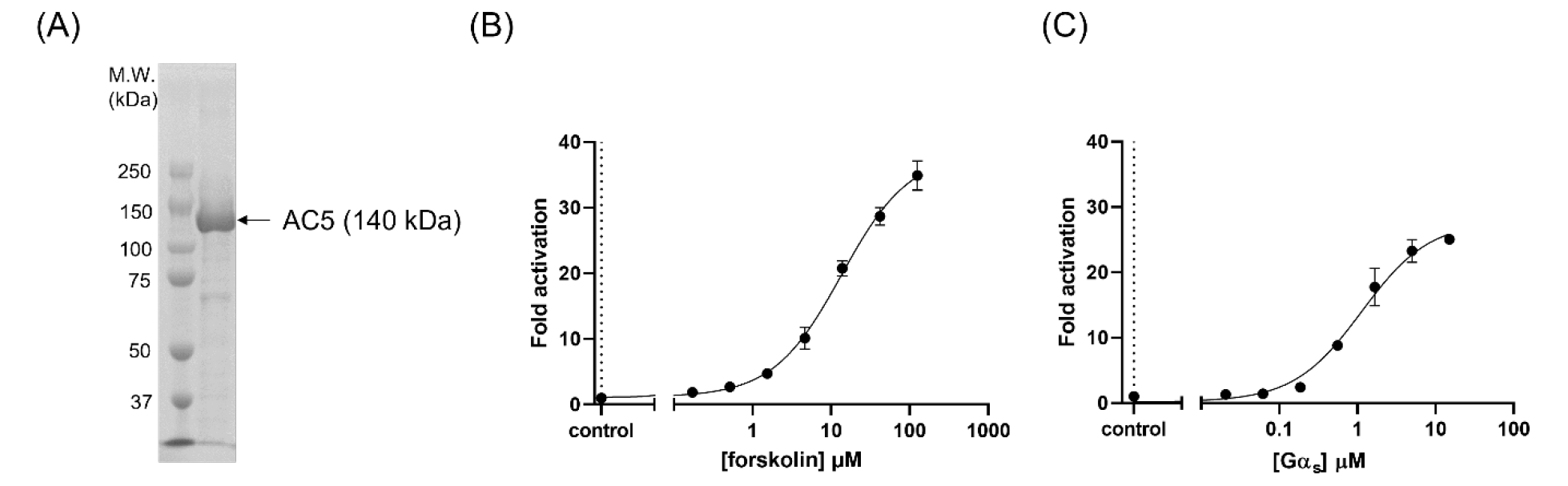
Purification and characterization of AC5. (A) AC5 was purified using anti-FLAG M2 affinity resin and analyzed by Coomassie blue stained SDS-PAGE. The AC activity of purified AC5 was determined in the presence of 10 mM MgCl_2_, 0.5 mM ATP at the indicated concentrations of (B) forskolin or (C) Gα_s_·GTPγS. The data shown are the mean ± SD (n=3).

**Figure S2.**
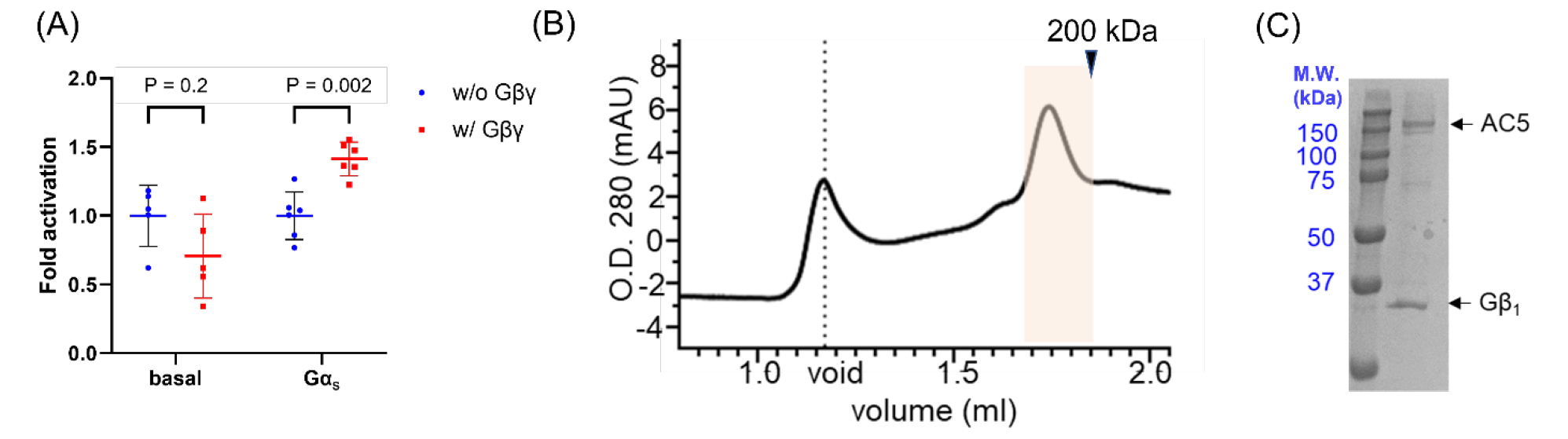
Activation of AC5 by Gβγ and purification of an AC5–Gβγ complex. (A) AC activity of purified AC5 was determined in the presence of 10 mM MgCl_2_ and 0.5 mM ATP with or without 250 nM Gα_s_·GTPγS, in the absence (•) or presence (▪) of 400 nM Gβ_1_γ_2_. Data presented are the mean ± SD. (B) The AC5–Gβγ complex was purified using anti-FLAG M2 affinity resin and injected onto a Superose 6 Increase 3.2/300 column equilibrated with 50 mM HEPES pH 8, 100 mM NaCl, 10 mM MgCl_2_, and 0.01 % LMNG. Peak fractions highlighted in pink were pooled, concentrated, and (C) analyzed by SDS-PAGE (note that Gγ_2_ runs off the gel).

**Figure S3.**
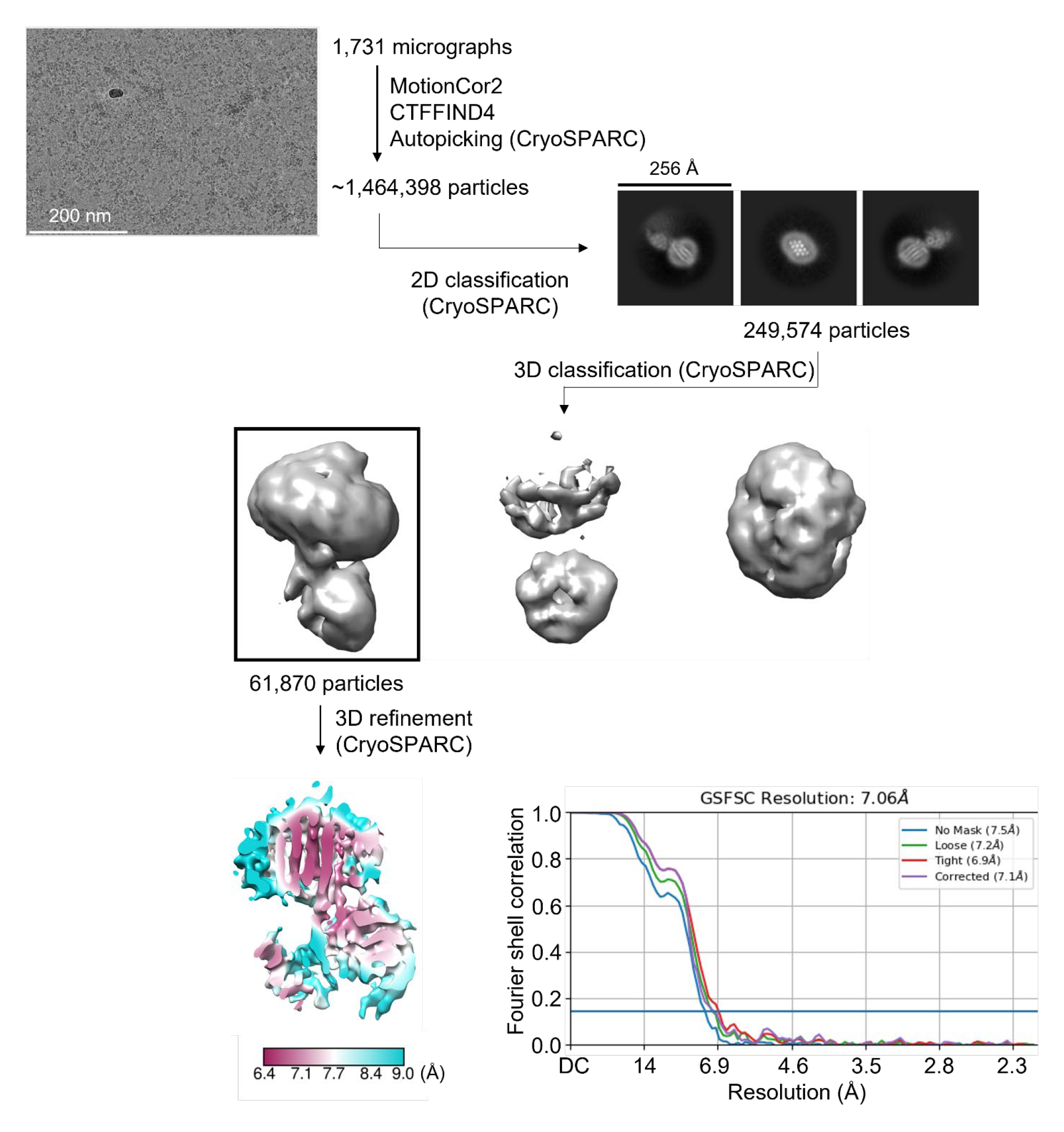
Cryo-EM data processing workflow and resolution analysis of the AC5– Gβγ complex. The workflow, including a representative micrograph, 2D class averages (box size = 256 Å, mask = 180 Å), local resolution map, and Fourier shell correlation (FSC) curves calculated from two independent reconstructions by CryoSPARC. The nominal resolution of the resulting map, as defined by the 0.143 cutoff, is indicated by the horizontal blue line.

**Figure S4.**
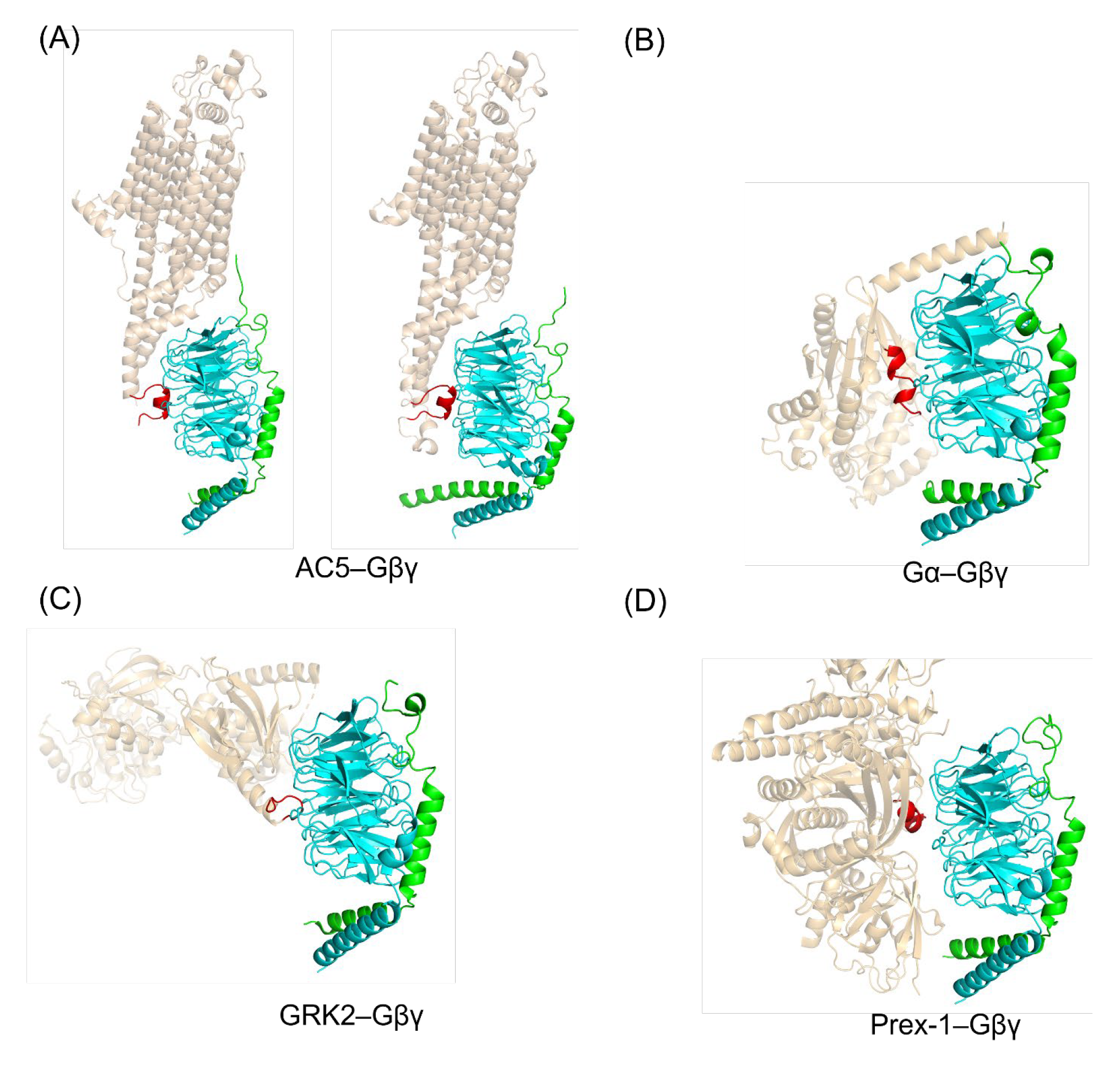
Binding mode of the AC5–Gβγ complex predicted by AlphaFold Multimer^19^ and comparison with other Gβγ complexes. (A) The docked model of AC5–Gβγ complex determined in this study (left, PDB entry 8SL3) and that predicted by AlphaFold Multimer (right). For comparison, some other Gβγ complexes are shown, including (B) Gα (PDB entry 1GOT), (C) GRK2 (PDB entry 6U7C), and (D) Prex-1 (PDB entry 6PCV) in complex with Gβγ. The red highlighted region in each structure contacts the central core “hotspot” of Gβ close to or centered at Trp99.

**Figure S5.**
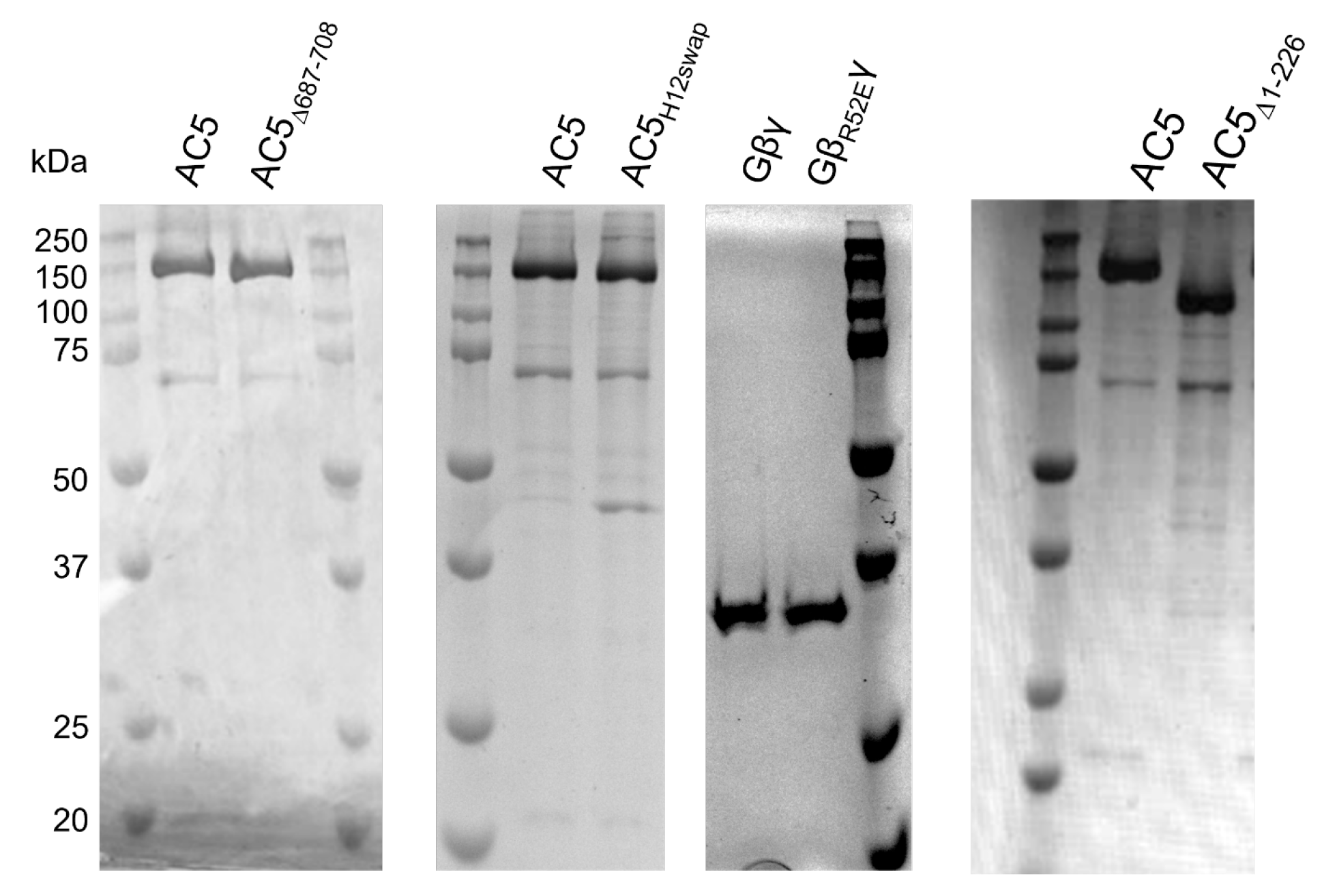
Purified AC5 and Gβγ variants used in this study. The purity of AC5 and Gβγ variants was assessed using Coomassie blue stained SDS-PAGE (Gγ at 7.5 kDa runs off the gel).

**Figure S6.**
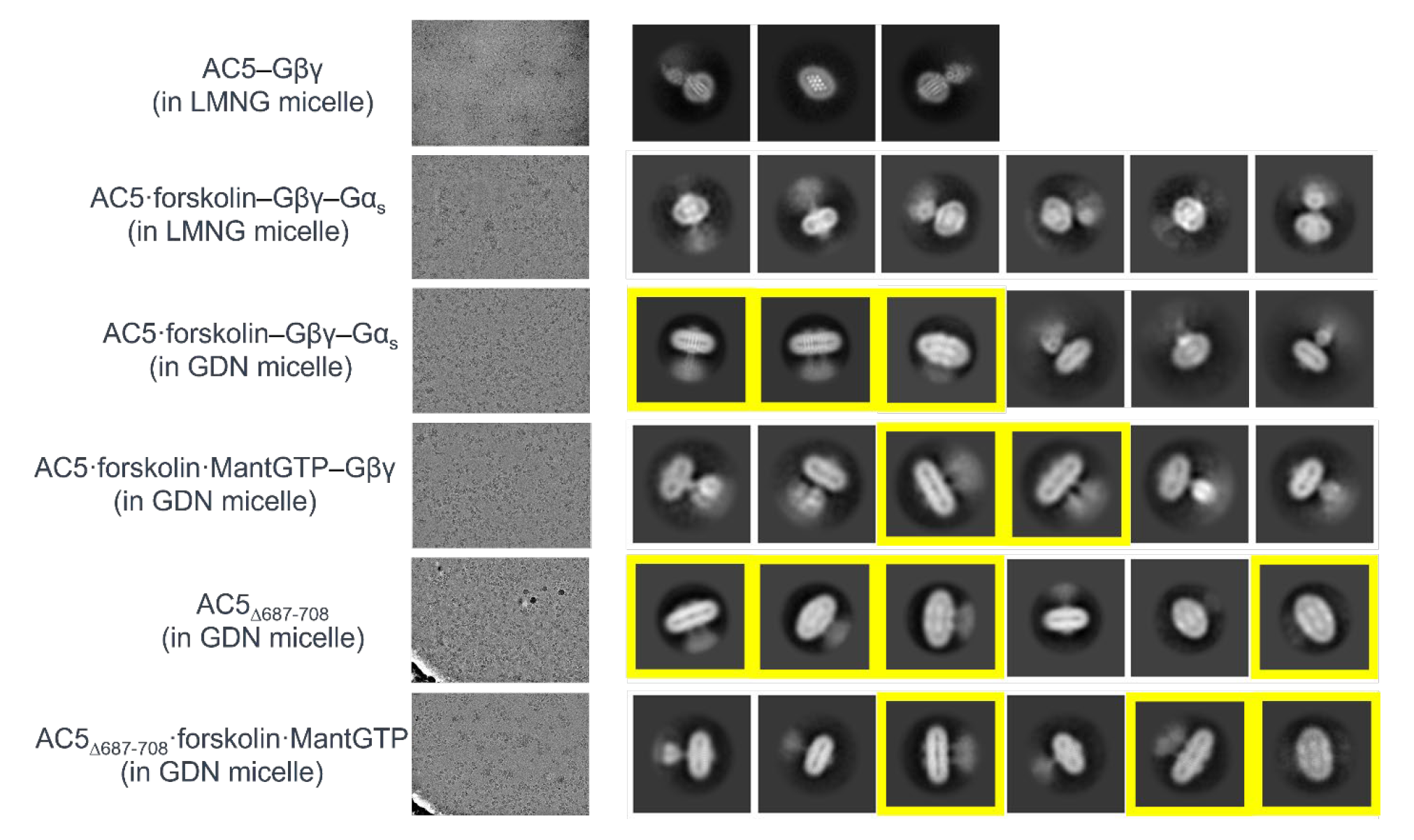
Cryo-EM analysis of various AC5 samples described in this manuscript. Representative micrographs and 2D class averages are shown for each sample. Note that homodimers (yellow boxes) are only observed in GDN regardless of activation status.

**Figure S7.**
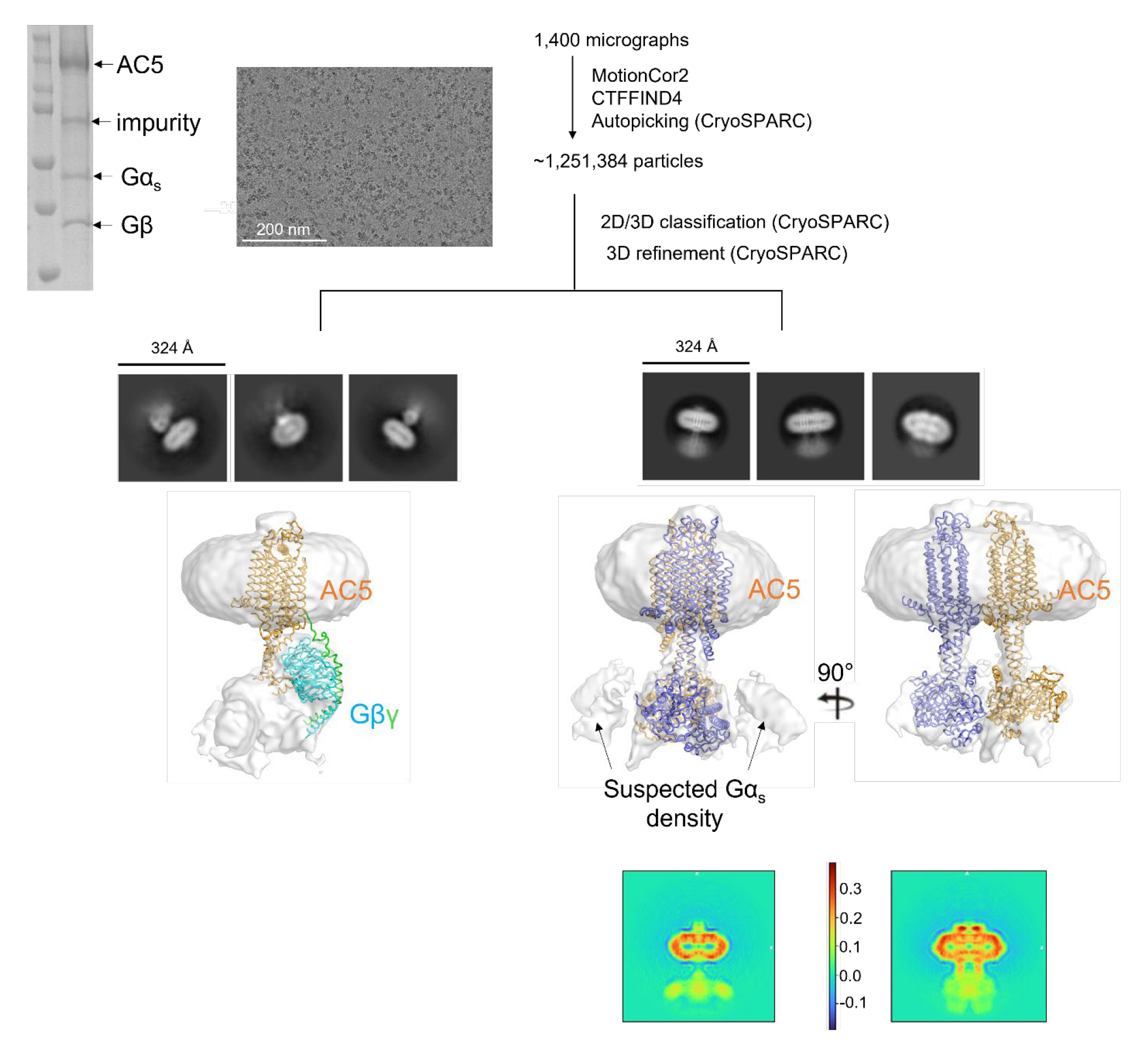
Sample preparation and cryo-EM data processing workflow of the AC5– Gβγ–Gα_s_ complex. A presumptive AC5–Gβγ–Gα_s_ complex was purified using Anti-FLAG antibody pulldown and analyzed by SDS-PAGE with Coomassie blue staining (top left). Cryo-EM data processing workflow includes a representative micrograph, 2D class averages (box size = 324 Å, mask = 200 Å for AC5–Gβγ on the left and 180 Å for dimeric AC5 on the right), and reconstructed 3D models. The dataset reveals two distinct populations, with one corresponding to the AC5–Gβγ monomer (bottom left) and the other an AC5 dimer containing density corresponding to Gα_s_ subunits (bottom right). Below the dimeric AC5 map, the central slice is displayed in two directions. Note that the 3×4 array of TM helices line up in the slice on the right into three vertical lines in each subunit, consistent with the above cartoon. The scale bar of the heat map indicates arbitrary units of density obtained from the particle images.

**Figure S8.**
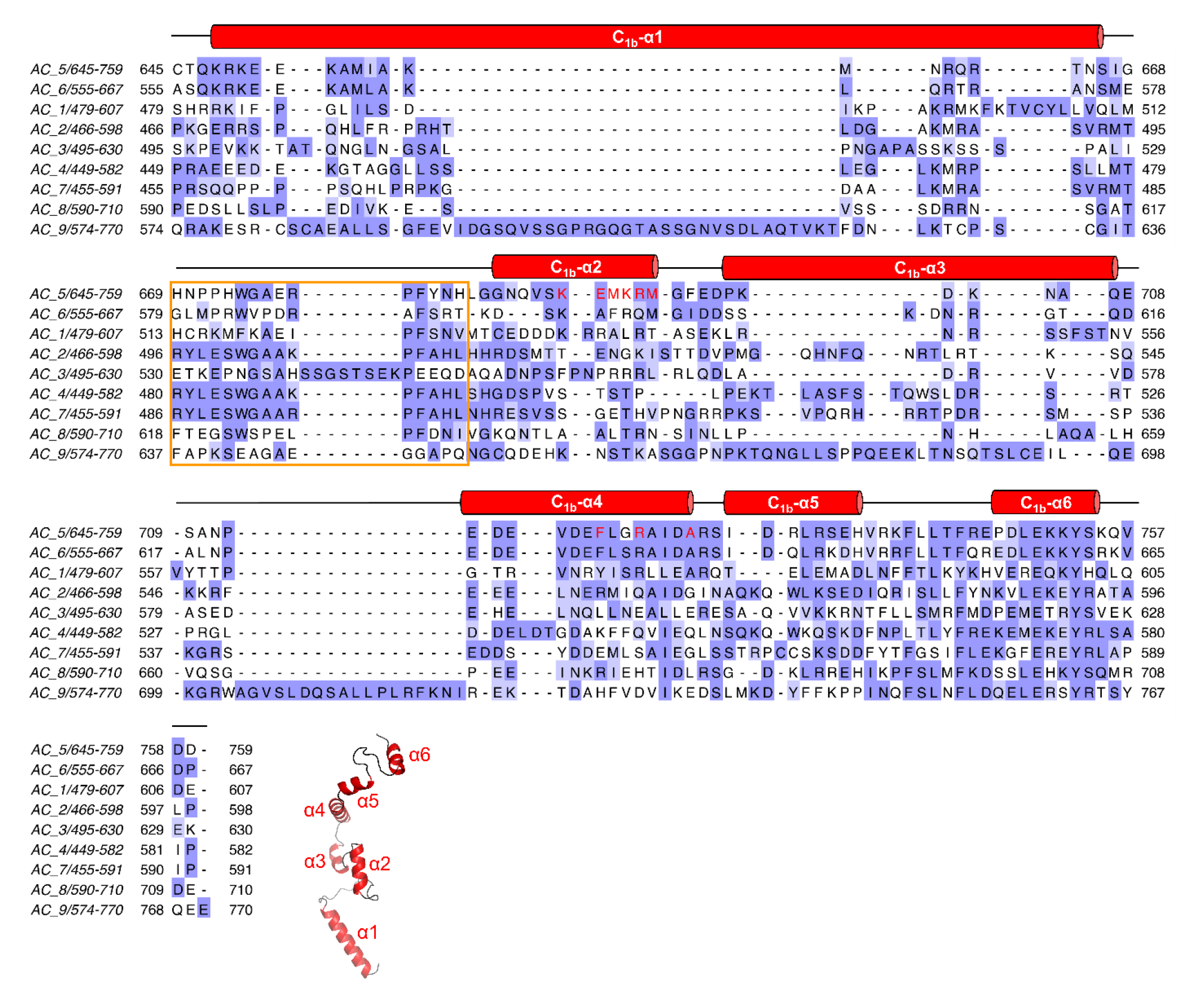
Sequence alignment of the C_1b_ region. Residue similarities are colored according to the BLOSUM62 score. Secondary structures of the primary sequence for the AC5 structure here are displayed on top, and the AlphaFold2 predicted model for the C_1b_ region in AC5 is shown at the bottom as a reference for the named helices. The predicted “PFAHL” Gβγ binding site in AC2 is highlighted in an orange box. Residues modelled in contact with Gβγ in the AC5–Gβγ complex (PDB entry 8SL3) are drawn in red color.

**Figure S9.**
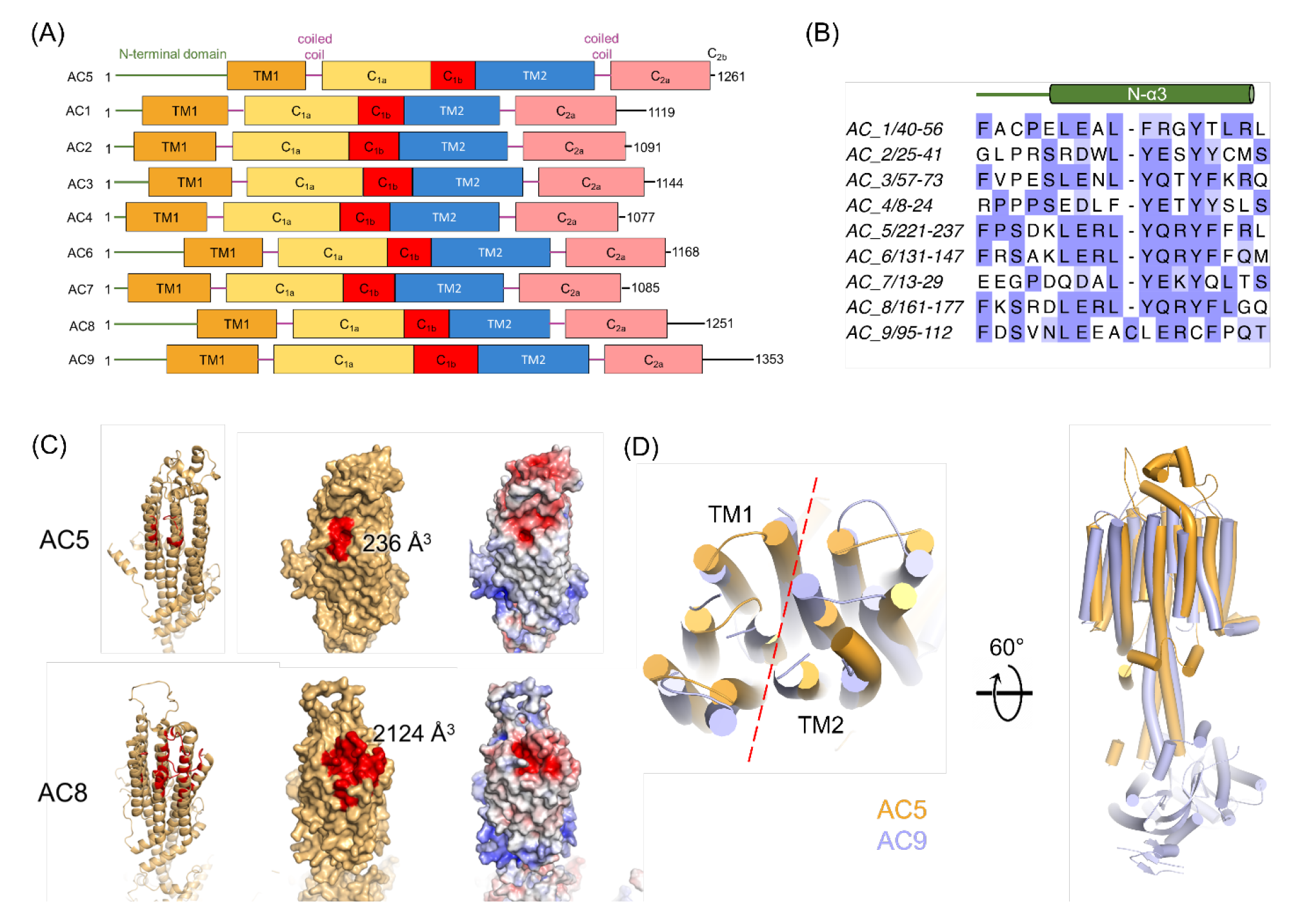
Comparison of mAC structural features. (A) Domain architecture of mAC isoforms. (B) Sequence alignment of the N-α3 helix among mAC isoforms, with secondary structure from AC5 indicated above. (C) Comparison of a predicted binding pocket in the extracellular domain of AC5 and AC8, with the residues predicted to form the binding pocket highlighted in red in both the cartoon (left) and surface (middle) representations. Additionally, the electrostatic potential is shown (right), highlighting the negatively charged surface of the extracellular domain. (D) Superposition of AC5 (PDB entry 8SL3) and AC9 (PDB entry 6R3Q), revealing similar organization of the 12 TM helices (with the extracellular domain of AC5 removed for clarity, left) but subtle differences in the coiled-coil domain (right).

**Table S1.** EC_50_ values of Gβγ for AC5 variants.

| Purified protein ( in GDN micelle) |  |  | Membrane |  |  |
| --- | --- | --- | --- | --- | --- |
| Variants | log EC<sub>50</sub> (nM) | EC<sub>50</sub> (nM) | Variants | log EC<sub>50</sub> (nM) | EC<sub>50</sub> (nM) |
| AC5 | 2.7 ± 0.1 | 460 | AC5 | 1.6 ± 0.2 | 40 |
| AC5<sub>Δ687-708</sub> | 3.5 ± 0.2 | 3100 * | AC5<sub>Δ687-708</sub> | 2.4 ± 0.3 | 270 * |
| AC5<sub>Δ1-226</sub> | 3.0 ± 0.1 | 970 | AC5<sub>R418W</sub> | 2.4 ± 0.5 | 250 * |
| AC5<sub>H12swap</sub> | 3.0 ± 0.2 | 1000 * | AC5<sub>Δ9bp</sub> | 2.8 ± 0.4 | 300 * |
|  |  |  | AC5<sub>M1029K</sub> | 2.5 ± 0.2 | 630 |

\*Curve did not reach saturation, thus the EC<sub>50</sub> value is an estimate.

**Table S2.** Cryo-EM data collection, refinement, and validation statistics.

| Sample: | AC5-Gβy<br>(EMDB-40572)<br>(PDB 8SL3) | Dimeric AC5<br>(EMDB-40573)<br>(PDB 8SL4) |
| --- | --- | --- |
| **Data collection and processing** |  |  |
| Magnification | 81,000 X | 81,000 X |
| Voltage (kV) | 300 | 300 |
| Electron exposure (e-/Å<sup>2</sup>) | 53.8 | 53.8 |
| Defocus range (μm) | 1 ~ 3 | 1 ~ 3 |
| Pixel size (Å) | 1.08 | 1.08 |
| Symmetry imposed | C1 | C1 |
| Initial particle images (no.) | 1,464,398 | 1,251,384 |
| Final particle images (no.) | 61,870 | 102,581 |
| Map resolution (Å) |  |  |
| FSC threshold 0.143 | 7.0 | 6.6 |
| **Refinement** |  |  |
| Map sharpening *B* factor (Å<sup>2</sup>) | 695 | 343 |
| Map CC | 0.47 | 0.34 |
| Model composition |  |  |
| Non-hydrogen atoms | 7,675 | 15268 |
| Protein residues | 977 | 1940 |
| Ligands | CM1:1 | NA |
| *B* factors (Å<sup>2</sup>) |  |  |
| Protein | 43 | 38 |
| Ligand | 16 | NA |
| R.m.s. deviations |  |  |
| Bond lengths (Å) | 0.004 | 0.003 |
| Bond angles (°) | 0.81 | 0.647 |
| Validation |  |  |
| MolProbity score | 2.02 | 1.56 |
| Clashscore | 10.55 | 8.12 |
| Poor rotamers (%) | 1.9 | 1.01 |
| Ramachandran plot |  |  |
| Favored (%) | 96.07 | 97.40 |
| Allowed (%) | 3.10 | 2.18 |
| Disallowed (%) | 0.83 | 0.42 |

**Table S3.**
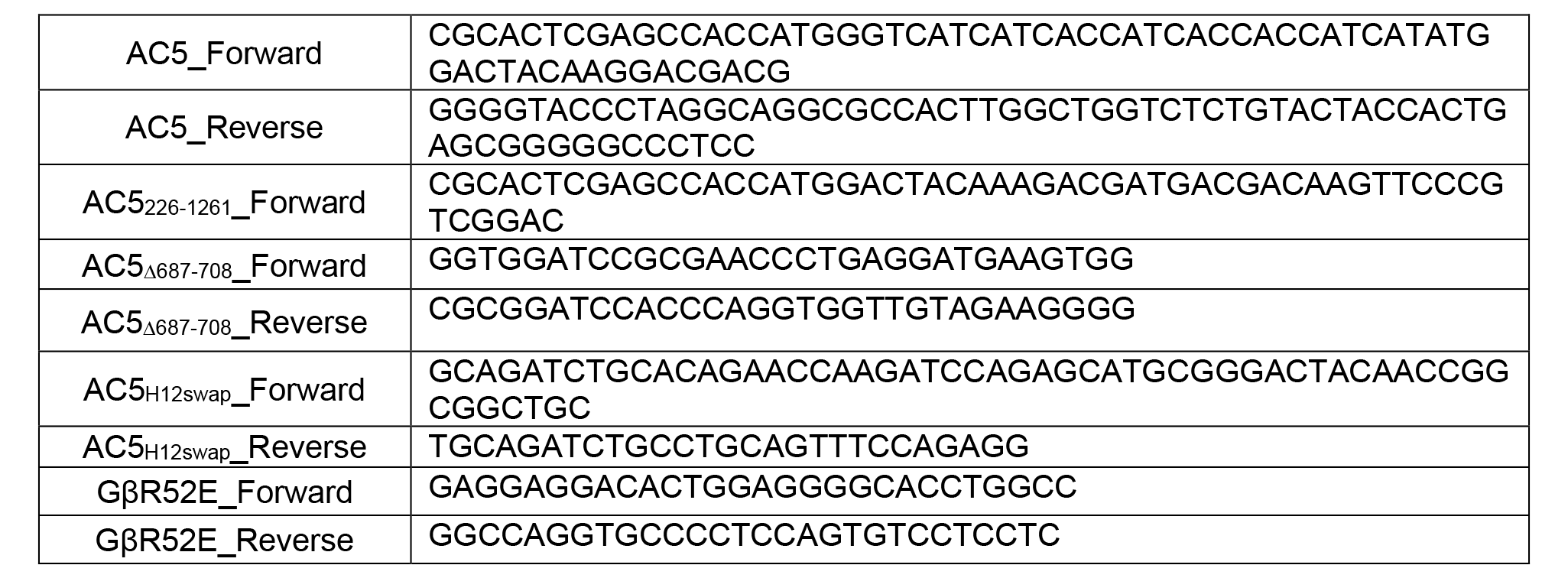
List of primers used.

## Notes

### Competing Interest Statement

The authors have declared no competing interest.

